# Developmental trajectory of thalamus topography during the late preterm and perinatal period in normal development, after premature birth and in congenital heart defects

**DOI:** 10.1101/2024.08.27.608544

**Authors:** Hui Ji, Zhexin Wu, Anna Speckert, Ruth Tuura, Cornelia Hagmann, Beatrice Latal, Walter Knirsch, Andras Jakab

## Abstract

The thalamus plays a critical role in neural circuit maturation and brain connectivity. It is well known to show a characteristic topographical organization of local and long-range connectivity, yet developmental changes in this topography remains incompletely understood. Our study aims to first validate the use of diffusion MRI for the topographic delineation of thalamic subdivisions (nuclei), to characterize thalamic topography during development and assess the impact of congenital heart defects (CHD) on thalamus development.

We used local fiber orientation distribution functions, derived from diffusion MRI (dMRI) data, to subdivide the thalamus into seven clusters. We first evaluated the within-subject reproducibility of a dMRI based clustering method across 30 adults scan-rescan datasets (n = 30) using K-means clustering, achieving reproducible segmentation of the thalamus into seven parts. This was applied to a large developmental cohort comprising normally developing infants from the term-born dHCP cohort (n = 68, gestational age [GW] at MRI scan ranging from 37.4 to 44.8 weeks, mean ± SD: 41.4 ± 2.1 weeks) and the Zurich cohort (n = 42, GW at MRI ranging from 39.0 to 48.7 weeks, mean ± SD: 42.55 ± 1.96 weeks). The cohort also included preterm-born dHCP cohort (n = 48, GW at MRI scan ranging from 29.3 to 37.4 corrected gestational weeks, mean ± SD: 34.4 ± 2.02 weeks) and one case of a postmortem fetal specimen (n = 1, GW at MRI = 21 weeks). Cluster volumes were analyzed as a function of age using multivariate linear regression. Differences in cluster volumes between control subjects (n = 42, GW at MRI ranging from 39.0 to 48.7 weeks, mean ± SD: 42.55 ± 1.96 weeks) and newborns with congenital heart disease (CHD) (n = 49, GW at MRI ranging from 37.6 to 42.9 weeks, mean ± SD: 40.23 ± 1.30 weeks) were analyzed using variance analysis, controlling for sex and age at MRI scan.

The clustering on average had a within-subject overlap of 0.811 (Dice overlap metric). While absolute thalamus volumes increased with age, their relative volumes remained stable in the studied developmental period. CHD infants shown reduced absolute volumes in six of the seven thalamic nuclei, with significant increasing in the relative volume of the cluster overlapping with the mediodorsal nucleus.

While the perinatal period comprises rapid developmental events including the maturation of white matter, the topography of thalamic circuitry remains stable. Our study highlights alterations in thalamic topology and potential impacts on brain connectivity in CHD infants. By leveraging local diffusion properties from diffusion MRI, our validated segmentation technique offers a robust, data-centric approach to thalamic delineation.

## 1. Introduction

The thalamus is essential for neural circuit development, driving key neurodevelopmental processes such as axonal guidance and neuronal migration (Miller et al. 1993; Auladell et al. 2000; Sur and Rubenstein 2005; Jones 2007). It plays a critical role in brain maturation and connectivity not only as a sensory signal relay station but also by modulating information exchange between cortical regions. Its functional roles include sensorimotor, cognitive, emotional, and attentional functions. Previous studies have highlighted the importance of the thalamus, showing that disruptions are linked to various developmental or psychiatric conditions, including bipolar, autism, cognitive, and emotional dysfunctions (Nair et al.; Carrera and Bogousslavsky 2006; Kubota et al. 2013; Alcauter et al. 2014; Anticevic et al. 2014; Nosarti et al. 2014). Disruptions in thalamic development and topographical organization have also been observed in preterm born infants and individuals with congenital heart defects (CHDs) (Ball et al. 2012; Ball et al. 2013; Ball et al. 2015; Jaimes et al. 2018). During early brain development, the thalamus establishes its topological structure, providing scaffolding for neural pathways, shaping brain function integration, and forming reciprocal connections with the cortex that significantly influence cortical development. Despite its importance, the emergence and reshaping of the thalamic connectional topology during the perinatal period remain poorly understood. This period involves increased sensory load as infants experience postnatal life. Extrapolating information from other species is challenging due to unique aspects of human brain development, such as the abundance of prefrontal connections and neuron counts in the mediodorsal nucleus (Abitz et al. 2007). Furthermore, the precise organization of early thalamic topographical structure, particularly the development and organization of thalamic nuclei during perinatal period, is not well known. Additionally, the impact of congenital disorders such as preterm birth and CHD on thalamic topographic structure remains poorly understood.

The thalamus, essential for numerous brain functions, contained specialized nuclei that are particularly vulnerable to specific neurological disorders. In neonates, thalamic circuitry development is particularly sensitive. Preterm infants often displaying reduced thalamic volume, indicating the impact of premature birth on the thalamus and overall brain architecture (Thompson et al.; Ball et al. 2012; Nosarti et al. 2014; Alexander et al. 2018; Menegaux et al. 2021). Moreover, neonates with congenital heart disease (CHD) exhibit varying degree of thalamic dysmaturation, ranging from normal development to significantly disorganization or reduction in thalamic nuclei (Jaimes et al. 2018). Young individuals with CHD also showed morphological alterations in thalamic structure (Brossard-Racine Gilbert et al. 2020).

Delineating thalamic nuclei is traditionally performed using histological methods and manual segmentation. However, advancements in magnetic resonance imaging (MRI) have enabled the *in vivo* parcellation, revealing detailed anatomy and connectivity (Eugenio Iglesias et al. 2018; Su et al. 2019). In recent years, functional MRI (fMRI) and diffusion MRI (dMRI) have been increasingly used to incorporate patient-specific information. While fMRI maps neural activity and reveal strong functional specialization, it does not align well with the histologic atlases (Iglehart et al. 2020). Diffusion tensor imaging (DTI) has become a commonly studied approach for thalamus parcellation, revealing both global and local brain connectivity (Wiegell et al. 2003; Ziyan et al. 2006; Jonasson et al. 2007; Jbabdi et al. 2009; Takahata et al. 2010; Ye et al. 2013). Connectivity-based parcellation assumes that the thalamic nuclei have characteristic connectivity profiles. Data-driven approaches have been used to subdivide the thalamus into regions with localized connectional or similar tissue microstructure properties (Behrens et al. 2003; Kumar et al. 2015; Battistella et al. 2017). In the developing human brain, pre-myelinated axons reduce diffusion anisotropy, thereby could diminishing the accuracy of long-range connectivity predictions (Dubois et al. 2014). DTI modelling dMRI signal at a local scale, based on principal diffusion orientations, can identify thalamic subdivisions but may result in spatially disconnected regions due to its focus on fiber directions (Unrath et al. 2008; Mang et al. 2012; Kumar et al. 2015). Recent advances have integrated spatial and diffusion tensor similarities, improving thalamic parcellation accuracy (Battistella et al. 2017). Further enhancements involve modeling fiber tract geometries using spherical harmonics (SH) basis to provide a comprehensive angular diffusion profile at each voxel, achieving more consistent DTI-based thalamic parcellation. The constrained spherical deconvolution (CSD) approach, particularly the single-shell 3-tissue CSD (SS3T-CSD) model, refines white matter fiber orientation distributions (FODs) by also considering grey matter and cerebral spinal fluid (CSF) contributions. This method might be suited to the neonatal brain’s distinct microstructure, characterized by higher free water content (Dhollander and Connelly 2016; Dhollander et al. 2019).

Our study aims to describe the developmental trajectory of the topographical organization of thalamic local diffusion properties and short-range connectivity. We use data from infants with congenital heart defects and after premature birth as a possible model for disrupted topographical organization. Building on previous work validating the use of local diffusion features for thalamic parcellation, we hypothesize that data driven clustering of the thalamus into sub-compartments based on the white matter FODs sufficiently reflect thalamic topography in early postnatal development. We optimize the clustering techniques to create anatomically meaningful subdivisions using two computational methods. Our code for clustering is available at https://github.com/10258392511/Clustering.

## 2. Methods

The first part of our study focused on developing and optimizing a thalamus clustering pipeline that utilizes fiber orientation distributions (FODs) derived from diffusion MRI. We employed a scan-rescan dataset of healthy adults, referred to as **Dataset A**, for this purpose. This approach was chosen because histological thalamus atlases are only available for adult brains, which typically provide higher quality scans and the availability of scan-rescan data allows for reliable evaluation criteria. For characterizing the development of thalamic topography in normal and pathological conditions, we utilized further datasets, described as **Datasets B-E** below.

In **Chapter 2.3** and **Chapter 2.6**, we utilized grid search and Bayesian optimization for feature engineering on scan-rescan data (**Dataset A**). In **Chapter 2.7**, we evaluated clustering accuracy using the Dice similarity score across 30 subjects from **Dataset A**. After optimizing the clustering parameters and achieving good accuracy, we employed K-means clustering and Gaussian Mixture Models (GMM) with the optimized parameters to **Datasets B-E**.

**Dataset B**, part of the Developing Human Connectome Project (dHCP), includes data from preterm and term-born infants. Together with Dataset C, which consists of Zurich newborn data, our objectives were to establish an overall growth trajectory of the thalamus by pooling data, examine the impact of different scan centers on segmentation results, and determine if preterm birth affects thalamic volume and topography. **Dataset D** contains data on congenital heart defects (CHD) from Zurich’s Heart and Brain study. By comparing it with **Dataset C**, we aimed to investigate the impact of CHD on thalamic development and structural topography. Lastly, **Dataset E** includes postmortem data with a gestational age of 21 weeks. We aimed to assess if our method is applicable to postmortem data and to explore differences in thalamic organization between earlier developmental stages and those of preterm and term-born newborns.

### 2.1. Study population and MRI acquisition

#### Dataset A (test-retest dataset): demographics and MRI acquisition

Dataset A is a validation dataset consisting of health adult subjects. A description of this open source dataset was previously published (Boekel et al. 2016). The dMRI datasets were retrieved from the test-retest repository, accessible at: http://www.nitrc.org/projects/dwi_test-retest/.

A total of 34 healthy young subjects underwent MRI scans twice within the same day using a 3T Philips Achieva XT scanner equipped with a 32-channel head coil. However, Post-processing errors led to the exclusion of four subjects from the analysis. Specifically, one subject was excluded due to a significant discrepancy in slice thickness between initial and repeated scans. Another was excluded due to inadequate registration from T1-weighted MRI to diffusion MRI, and the remaining two were excluded due to registration errors between the thalamus atlas and individual thalamic structures. Consequently, the final cohort comprised 30 participants. T1-weighted anatomical scan was acquired using a T1 turbo field echo sequence. The parameters of this sequence were as follows: TR/TE = 8.4/3.9 ms, flip angle = 8°, voxel size = 1 × 1 × 1 mm^3^, and 220 coronal slices. For diffusion MRI scan, each subject underwent two scanning session in a single day. In each session, the diffusion MRI was repeated four times using a multi-slice spin echo (MS-SE), single-shot DWI scans with TR/TE = 7.545/86 ms. voxel size = 2 × 2 × 2 mm^3^, 60 transverse slices, and diffusion weighting along 32 gradient directions (b value = 1000 s/mm^2^). Additionally, six non-diffusion weighted images (b0; b value = 0 s/mm^2^) were acquired and subsequently averaged. To minimize motion artifacts, only the first diffusion MRI scan from each session was used in the subsequent analyses.

#### Dataset A: MRI processing

The thalamus proper (= thalamus outlines excluding the LGN and MGN) was manually annotated using 3D Slicer (Kikinis et al. 2014), version 4.8, on the 3DT1 images. To accurately transform the structural thalamus mask to diffusion space, our methodology was structured with following steps. Initially, we extracted the b=1000 s/mm^2^ images from diffusion MRI data and calculated their median using *dwiextract* in MRtrix3 (Tournier et al. 2019) and *fslmaths*. Following this, bias field correction was applied to the median b=1000 s/mm^2^ image utilizing *dwibiasfield* correction in ANTs (Tustison et al. 2010). We then aligned the 3DT1 images to the bias field corrected median b=1000 s/mm^2^ images via linear registration using FLIRT in FSL with 6 degrees of freedom (DOF) (Jenkinson and Smith 2001). The linear transformation matrix obtained was converted into an ANTs compatible format using the c3d_affine_tool from the c3d software. Next, 3DT1 images were registered to the median b=1000 s/mm^2^ image, both linearly and non-linearly, using the antsRegistrationSyN script in the ANTs toolbox (Avants et al. 2008) and the previously estimated linear transformation matrix serving as the initialization step. Finally, the thalamus masks were transformed from the 3DT1 space to the subjects’ diffusion space using the resulting transformations using antsApplyTransform, and then were thresholded at 0.5 in the diffusion space to rectify errors in the linear interpolation.

#### Dataset B (developing Human Connectome Project data): demographics and MRI acquisition

Dataset B was used to describe the development of the thalamic topography in normal newborns and in prematurely born babies. This dataset was sampled from the open-source developing Human Connectome Project (Bastiani et al. 2018), neonatal release 2, accessed 1/2022. For our study, we sampled the dHCP release 2 for a balanced number of subjects for each gestational weeks at scan time, including preterm born subjects as well as term born infants. The sampled dataset included MRI and demographic data (birth age, scan age, sex) for 120 subjects. After excluding four cases due to unavailable diffusion MRI data, a total of 116 cases (56 females and 60 males) were analyzed. The birth age range was 25.6 to 42 weeks (mean ± SD, 36.8 ± 4.3), and the scan age range from 29.3 to 44.7 (mean ± SD, 38.5 ± 4.02) weeks. The cohort included 48 preterm born infants with birth age ranging from 25.6 to 36.9 (mean ± SD, 32.5 ± 3.1) weeks and scan age ranging from 29.3 to 37.4 (mean ± SD, 34.4 ± 2.02) weeks. Additionally, 68 term born infants were included, with birth ages ranging from 37 to 41.9 (mean ± SD, 39.9 ± 1.3) weeks and scan age ranging from 37.4 to 44.8 (mean ± SD, 41.4 ± 2.1) weeks.

Anatomical MRI: T2-weighted images were acquired with a Turbo Spin Echo (TSE) sequence, were processed using super-resolution reconstruction (Kuklisova-Murgasova et al. 2012) and motion correction (Cordero-Grande et al.), producing 3D super-resolution reconstructed T2 volume brain (further referred to as 3DT2) with isotropic resolution of 0.5 × 0.5 × 0.5 mm^3^. we employed the 3DT2 from the dHCP open database. The MRI acquisition and parameters further detailed (Makropoulos et al. 2018). Diffusion MRI: The diffusion protocol utilized a spherically optimized scheme across four b-values (b0: 20; b=400 s/mm^2^: 64; b=1000 s/mm^2^: 88; b=2600 s/mm^2^: 128), ensuring comprehensive angular resolution. Scans were conducted at a resolution of 1.5 × 1.5 mm with 3 mm slice thickness and imaging parameters of TR/TE = 3800/90 ms. Additional diffusion MRI protocol details can be found in publications (Bastiani et al. 2018).

#### Dataset B: MRI processing

In Dataset B, the structural MRI data were sourced from the Developing Human Connectome Project (dHCP) and had been pre-processed using the dHCP structural MRI pipeline. This pipeline segmented the 3DT2 images into 87 anatomical labels, out of which we extracted the left and right thalamus masks separately. Similarly, the diffusion MRI data were also pre-processed via the dHCP diffusion MRI pipeline (Bastiani et al. 2018), incorporating motion correction and eddy current distortion corrections with techniques from FSL (Andersson and Sotiropoulos 2016). To further process the dMRI data for our specific study, we applied bias-field correction using MRtrix3 to correct for intensity inhomogeneity (Tustison et al. 2010). Subsequently, the 3DT2 thalamus mask were aligned to the diffusion space. This involved extracting the b0 image from the diffusion data and calculating its median, aligning the 3DT2 images to the median b0 volume using FLIRT in FSL (Jenkinson and Smith 2001) for linear registration with 6 DOF. After this, the same procedure was followed to align the thalamus masks with a linear and non-linear transformation, mirroring the procedure used in Dataset A.

#### Dataset C and Dataset D: demographics and MRI acquisition

Dataset C included healthy term-born infants, recruited from January 2011 to April 2019 from the Heart and Brain Research Group Zurich at the University Children’s Hospital Zurich. Inclusion criteria for Dataset C (control group) specified birth at >36 weeks of gestation with unremarkable postnatal adaptation. Dataset D comprised neonates with CHDs, recruited between December 2009 and March 2019. The study was approved by the Ethical Committee of the Canton Zurich, with parental consent obtained for all participants.

Both groups underwent brain MRIs on a 3.0T GE Signa MR750 scanner with an 8-channel head coil. Anatomical MRI sequence included a T2-weighted fast-spin-echo sequence in axial, coronal, and sagittal planes (TE/TR: 97/5900 ms, flip angle: 90°, slice thickness 2.5 mm, slice gap 0.2 mm). Further details about the imaging setting and MRI sequence parameters for this study have been reported previously (Bertholdt et al. 2013). Diffusion tensor imaging (DTI) acquired in the axial plane using a pulsed gradient spin-echo echo-planar imaging sequence (TR/TE 3950/90.5 ms, FOV 18 cm, matrix 128×128, slice thickness 3 mm) using 35 diffusion encoding gradient directions at b-value of 700 s/mm^2^ and four b = 0 images. MRI acquisition details for both datasets have been documented previous work (Feldmann et al. 2020).

After DTI quality check, the control group (Dataset C) comprised 42 infants (19 females, 23 males), birth age range from 37.7 to 41.6 (mean ± SD, 39.58 ±1.22), scan age range from 39 to 48.7 (mean± SD, 42.55 ± 1.96) weeks, respectively. The CHD group (Dataset D) included 49 neonates (36 females, 13 males), ages at birth range from 36.5 to 41.4 (mean ± SD, 39.38 ± 1.28), scan age range from 37.6 to 42.9 (mean ± SD, 40.23± 1.30) weeks, respectively.

#### Dataset C and D: MRI processing

Axial, coronal and sagittal T2-weighted MR images were reconstructed into a 3DT2 image with an isotropic resolution of 0.5 mm using the SVRTK method (Kuklisova-Murgasova et al. 2012). dMRI data were preprocessed using in-house developed pipeline, which included denoising (Veraart et al. 2016), motion and distortion correction (Andersson et al. 2003; Andersson and Sotiropoulos 2016), bias-field correction (Tustison et al. 2010) using MRtrix3, FSL, ANTs (https://github.com/annspe/connectome_pipeline). Thalamic masks on the 3DT2 images were manually annotated using 3D Slicer (Kikinis et al. 2014). For the alignment of the 3DT2 thalamus masks to the diffusion MRI space in Datasets C and D, we followed the same methodology for transforming as used for Dataset B.

#### Dataset E (Post mortem fetal MRI)

We included a prenatal post-mortem human brain MRI data accessible from developing Allen Brain Atlas at (Allen Institute for Brain Science): http://www.brainspan.org/ scanned at 21 gestational weeks using a 3T Siemens MR scanner (Siemens Medical Solutions, Erlangen, Germany) with a custom solenoid coil. The diffusion imaging was performed with diffusion-weighted steady-state free precession (DW-SSFP) parameters: TR = 24.5 ms, TE = 18.76 ms, 0.4 mm isotropic resolution, 150 Hz/px bandwidth, 160×160×160 matrix, 42 diffusion gradient directions, and four b0 volumes.

#### Dataset E: MRI processing

We used preprocessed data from Allen Institute, which included temperature drift corrections using MATLAB, and co-registration of non-diffusion-weighted volumes with FSL’s FLIRT to correct for b0 inhomogeneity and eddy-current distortions. For this study, we performed bias field correction using ANTs (Avants et al. 2011). Due to the high-resolution acquisition (0.4 mm isotropic), thalamus masks were manually annotated directly on the diffusion MRI images. Detailed sample information, MRI scan parameters, and processing methods can be found in the Developing Human Brain Imaging whitepaper at www.brainspan.org.

### 2.2. Fiber Orientation Distribution (FOD) estimation from dMRI

In our study, we utilized local diffusion features to delineate thalamic topography through a data-driven clustering method. We applied constrained spherical deconvolution (CSD) in MRtrix3 to estimate the white matter fiber orientation distributions (WM-FODs) from the dMRI signal in each voxel. Using a spherical harmonics (SH) basis of orders 0, 2, 4, 6, and 8, we represented the WM-FODs as vectors with dimensions 1, 6, 15, 28, and 45, respectively.

These high-dimensional feature vectors describe the complex fiber orientations within each voxel. The SH basis provided an angular characterization of the WM-FODs, as depicted in the accompanying glyph figure in **Figure 1**. This approach allowed us to use simple distance metrics to assess similarities in diffusion properties across WM-FODs, aligning with the methodology (Wassermann et al. 2008) for clustering similar fiber orientations.

**Figure 1.**
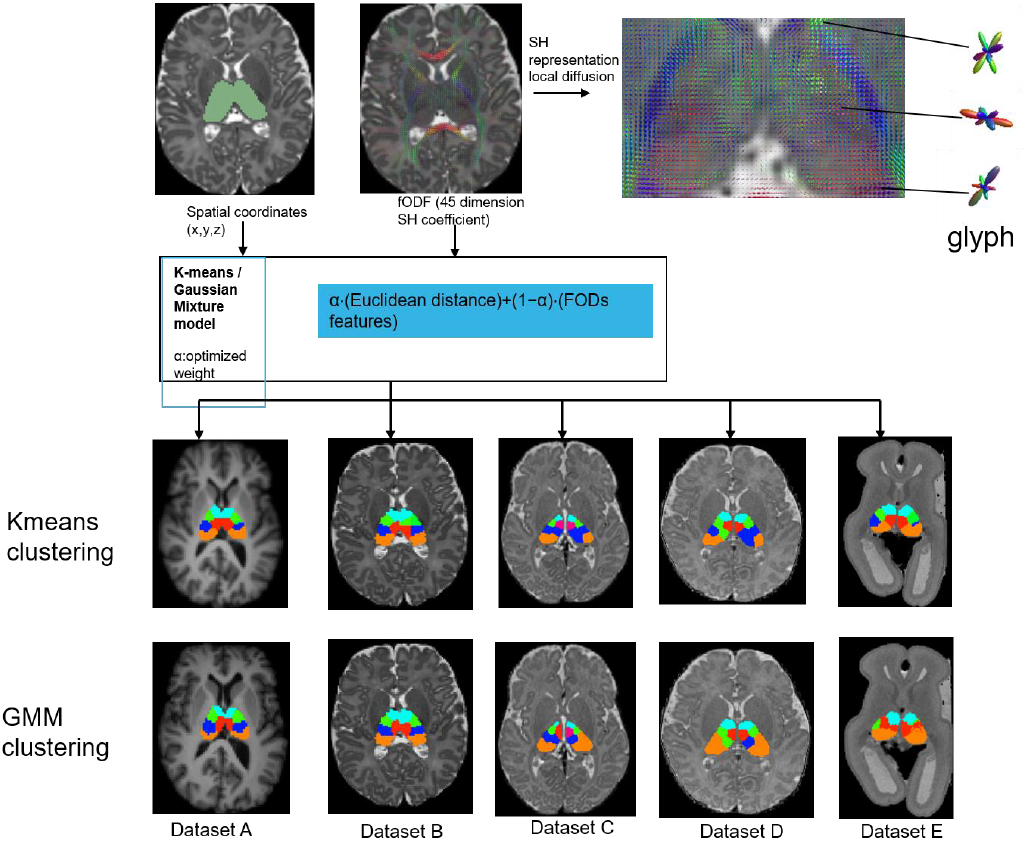
Overview of the clustering framework and clustering results, and an illustration of the WM-FOD as glyphs for the same MR image level. dMRI-based segmentation of the thalamic nuclei has been performed using a K-means and GMM clustering algorithm with the optimized features. The images show individual results of the thalamic nuclei segmentation and the glyphs corresponding to the WM-FOD in individual voxels. The bottom two rows illustrate the hard-clustering as well as probabilistic clustering (GMM) as overlay on the subject’s structural image for each study groups. Dataset A: adults subject, Dataset B: dHCP, Dataset C: normally developing newborn from the Zürich dataset, Dataset D: newborn with congenital heart defect, Data E: post mortem fetal MRI.

We first estimated white matter (WM), grey matter (GM), and cerebrospinal fluid (CSF) response functions for each subject using the *dhollander* algorithm using MRtrix3Tissue (https://3tissue.github.io), a fork of MRtrix3 (Tournier et al. 2004). Since the comparison of WM-FODs across subjects requires a group-average response function, we followed different procedures for each dataset. For dataset A (healthy adults for technical optimization), we utilized the single shell three tissue (SS3T) model (https://3tissue.github.io) in MRtrix3 with averaged response functions from all 30 subjects. For the Dataset B (dHCP), group-averaged response functions were derived by averaging the WM, GM, and CSF response functions of the 20 oldest subjects (GA 43-45 weeks), representing relatively mature white matter. For datasets C and D, we employed a uniform approach by using the average response functions from dataset C (normal controls) for both groups. This decision was driven by the need for consistency in group comparison, particularly since dataset D comprises subjects with congenital heart defects (CHD). Using the same unit response function from dataset C for both C and D ensures comparability.

Subsequently, each subject’s diffusion MRI signal was deconvolved into tissue FODs using the single shell three tissue CSD (SS3T-CSD) method, employing the respective group-average response functions. This approach enabled us to effectively use FODs as features for clustering, aiming to delineate distinct neural tracts based on their orientation and diffusion properties. Each subject’s resulting tissue FODs (i.e., WM-FOD, GM-FOD, CSF-FOD) underwent intensity normalization (Tournier et al. 2019). Given that the discrimination between thalamic nuclei is most likely possible based on the white matter architecture within the thalamus (e.g. bundles of axons), and since WM-FODs might the highest signal-to-noise ratio, we used the WM-FOD for our feature extraction.

### 2.3. Feature engineering for clustering

To cluster the thalamus into discrete subdivisions that may correspond to functional subunits or nuclei, an optimal selection of features was necessary. Our method used both WM-FODs and spatial coordinates derived from diffusion MRI data for voxel clustering, following the considerations of a previous work that revealed superiority of this approach compared to using long-range connectivity (Battistella et al. 2017).

For each voxel, we initially created two feature vectors, a spatial coordinate feature (x, y, z) and a WM-FODs feature comprising 45-dimensional spherical harmonic coefficients. These spatial and FOD-based features were scaled and concatenated into a fused feature vector. The combination was controlled by two hyperparameters: α (weight for spatial feature) and s (scale for spherical harmonic coefficients). The resulting combined feature vector for voxel ***i*** was a fusion of the scaled spatial and WM-FOD features.

To ensure consistency across differently MRI scans and subjects, spatial coordinates were normalized using two methods. First, spatial coordinates were normalized to lie within the range [-1, 1] while preserving the original aspect ratio of the MRI volume. Second, distances between voxel feature representations were computed in the fused feature space. This approach allowed us to effectively cluster thalamus voxels based on a combination of spatial location and diffusion characteristics, facilitating the identification of potential functional subunits or nuclei within the thalamus.

### 2.4. Clustering

To optimize the clustering hyperparameters α and s, evaluate within-subject reproducibility, and compare the results to histological/anatomical subdivisions of the thalamus, we applied K-means clustering to Dataset A. Based on cytoarchitectural studies, a possible way to group thalamic nuclei is into seven main groups (Morel et al. 1997). Previous diffusion MRI studies also support a cluster number of seven for the thalamus (Behrens et al. 2003). We used the Euclidean distance as the distance metric. After optimizing the hyperparameters, we applied the selected values to datasets B-E to characterize the trajectories and disease-related changes in the volumes of these subdivisions.

We employed two clustering methods, K-means clustering and Gaussian mixture models (GMM), using the *scikit-learn* package (Pedregosa et al. 2011). K**-**means clustering provided a hard assignment for each voxel, assigning it to the cluster with the nearest centroid based on the shortest Euclidean distance between the voxel’s features and the cluster centroids. In contrast to K-means, GMM allowed for soft label assignment, giving each voxel a continuous probabilistic distribution over the clusters. This approach is particularly advantageous for the thalamus in early brain development, where voxels may correspond to multiple functional groups, which can be effectively captured with probabilistic assignments.

For a set of observation ***X*** = {***x***_**1**_, ***x***_**2**_, **…**, ***x***_***N***_}, where each is a 48-dimensional vector, a GMM models the data as a mixture of *K* different Gaussian distributions. The likelihood of our data under this model is:

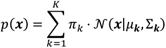

N(x|µ_k,_ ∑k)is the Gaussian distribution for cluster k with mean µ_k_ and covariance ∑_k_, and π_k_ represent the mixing coefficient for the *k-th* Gaussian (or π_k_ represent the mixing coefficient for cluster k), satisfying 0 < *π*_*k*_ < 1, and 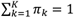.

### 2.5. Registration and atlas re-mapping

To evaluate the similarity of the thalamus clusters to anatomically (histologically) determined thalamic subdivisions, we utilized a publicly available thalamus atlas https://www.research-collection.ethz.ch/handle/20.500.11850/669887. Specifically, we used the MNI152-aligned, voxelized version of the Morel Atlas of the Human thalamus (Krauth et al. 2009; Jakab et al. 2012). This atlas was aligned to a neonatal T2-weighted imaging template from the ALBERTs atlas (Gousias et al. 2012) using the statistical shape model-based alignment method with the thalamus outlines serving as the shape predictor (Jakab et al. 2012).

The alignment of the thalamus outlines from the atlas to the subject’s thalamus outlines was performed in two registration steps. First, we conducted a linear registration where the T2-weighted template image was aligned to the subject’s T2 image using FLS’s FLIRT with 6 DOF. Second, another linear registration was performed, the thalamus atlas T2 image was then aligned using FLIRT with 9 DOF, with Y-axis rotations restricted to 20 degrees to improve the alignment of thalamus labels. This step used the thalamus outlines as guiding features rather than the grayscale images. Following these linear registrations, a non-linear outline-to-outline registration using the binarized label masks was performed using ANTs to further refine the alignment.

#### Atlas re-mapping

After aligning the atlas to the subject images, we re-numbered the K-means clustering-based thalamus clusters according to the label numbers corresponding to the closest atlas-based nuclei. The anatomical similarity between the clustering-based and subject-space histology-based subdivisions was assessed by calculating the overlap between the re-mapped atlas label images and the segmented thalamus clusters. This was quantified using the Dice similarity coefficient (DSC) and the 95th percentile Hausdorff distance (HD), as previously defined equations (Zou et al. 2004).

Given the Morel Atlas delineates over 25 anatomical nuclei, we grouped these into seven anatomically and functionally coherent groups to simply the analysis: anterior group (A), mediodorsal and central group (MDC), ventral anterior group (VA), ventrolateral and ventromedial group (VLM), ventral posterior group (VP), pulvinar and geniculate nuclei group (PuG), and lateral nuclei groups (L). Despite this grouping, we also report the overlaps of the clustering-based thalamus clusters with all individual Morel Atlas based labels as part of a qualitative evaluation of our clustering results.

### 2.6. Bayesian Optimization

We addressed the hyper-parameter tuning challenge by maximizing a function *g*: **R**^2^ → [0,1] where g links the hyperparameters *α* and *s* to the DSC value. To optimize this function, we employed Bayesian optimization with a Gaussian process (GP) as the surrogate model. The GP’s capability to model distributions over functions allows it to treat the function *g* as a sample from these distributions. This method enhances search efficiency and reduces computational expenses.

Optimization Model:

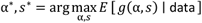

During the optimization process, each new observation *g* (α, s) refines the GP model, diminishing its uncertainty and focusing on promising region of the hyper-parameter space. Bayesian optimization leverages online learning to incrementally improve the estimate of a black-box objective function, making it particularly useful for problems where obtaining one observation is time-consuming. We used the Bayesian-optimization package for this implementation (Kim and Choi 2023).

### 2.7. Evaluation of clustering accuracy

The primary objective of optimizing clustering parameters was to enhance the specificity and sensitivity of the clustering procedure, ensuring that resulting clusters are both robust (consistent across repeat scans) and biologically valid (in agreement with histological findings). To assess the accuracy and robustness of our clustering method, we employed two primary strategies.

First, to evaluate reproducibility, we utilized the Dice Similarity Coefficient (DSC) to measure the overlap between initial scans and their corresponding re-scans from a test-retest dataset. For two scans, A and A’, and a thalamus subdivision C, the DSC for the scans across the 7 subdivisions is defined as:

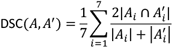

Where *A*_*i*_ and *A*_*i*_*’* represent the region of interest for the *i*-th sub-division in scans *A* and *A’*, respectively. We used the DSC implementation from MONAI (Jorge Cardoso et al. 2022). The DSC offers a quantitative measure of the spatial overlap between two samples and is computed for individual subjects as the mean DSC across the thalamus’ 7 distinct functional groups.

Second, to evaluate the biological relevance of our clustering method, we assessed its accuracy by comparing the clustering results with histological maps, which serve as a ground truth validation. By calculating the DSC between the clustering result and the histology-based atlas map, one can assess how well the clustering method identifies true anatomical or functional regions.

### 2.8. Selecting of the optimal number of clusters

To determine the optimal number of clusters for thalamus topography in developing cohorts, we employed three methods: the Elbow method, the Silhouette Score, and Gap statistics. The Elbow method identifies the point where additional clusters provide diminishing returns in explained variance. The Silhouette Score measures cluster cohesion and separation, with higher scores indicating better-defined clusters. Gap statistics compare clustering quality against randomly generated reference datasets to identify the most statistically significant clustering solution. Detailed descriptions of these methods and their application can be found in the **Supplementary Methods**, with the results of these experiments discussed in the **Supplementary Results**. Illustration of selecting optimal number of clusters are well described in **Supplementary Figure 1**

### 2.9. Statistical tests

In our study, thalamus topography is described as the developmental age dependent volumetric changes of the thalamus cluster volumes. We report absolute as well as relative volumes. To ensure the comparability across individuals and to describe topographical organization, each subdivision thalamus volumes were normalized to account for individual differences in overall thalamus size, denoted as relative volume.

We utilized robust regression to model the relationship between age and both absolute thalamic volume and normalized thalamus relative volume (Seabold and Perktold 2010). In our study, some of the clusters had zero values because two or more small clusters might have been re-mapped into the same cluster, we took the zero values as missing data in the statistical analysis. The model was fitted using the least square method.

To quantify age-related changes of thalamus volume, we carried out a linear regression analysis with age as the independent variable and absolute thalamic nuclei volumes and normalized relative thalamus nuclei volumes as the dependent variable. The significance of the regression coefficient for age determined if there’s a statistically significant age-related change in thalamus volume. This analysis was carried out by concatenating all developmental datasets together (Datasets B-D) and accounting for the dataset as a covariate in the model, as well as performing the test separately for each group, to account for possible differences in development between the groups.

To assess differences between the congenital heart disease (CHD) group and the control group, we utilized an ANOVA, correcting for sex and age at MRI. To account for multiple comparisons across the seven thalamic nuclei groups, we applied the False Discovery Rate (FDR) correction using the Benjamini-Hochberg method. All statistical analyses were conducted using the Python *StatsModels* package (Seabold and Perktold 2010).

## 3. Results

### 3.1. Parameters optimization

#### 3.1.1. Grid search

We adopted a grid search strategy aimed at optimizing the DSC between the within-subject repeated scans of Dataset A. As observed from Figure 1, both normalized spatial coordinates and the scaled SH coefficients independently contributed to achieving relatively high DSC values. This suggests that while spatial coordinates can serve as an effective feature for high DSC parcellation, SH coefficients also have the potential to provide comparable parcellation results. Our evaluation extended across various SH basis orders, specifically 0, 2, 4, 6, and 8, as illustrated in Figure 1.

#### 3.1.2. Bayesian Optimization

As a second step in optimizing the parameters for clustering, Bayesian optimization was employed after fixing the maximum spatial weights at 0.9, 0.8, 0.7, and 0.5. Our findings indicate that optimal DSC values for thalamus segmentation can be achieved using spatial weights significantly below 1. As presented in **Figure 2** and in **Supplementary Table 1**, we noticed two findings: firstly, high DSC values require a careful calibration of spatial weight and SH scaler; secondly, SH coefficients are crucial for thalamus clustering. While exact optimal values for spatial weight (α) and the spherical harmonic scaler (s) remain to be determined, our data indicate that the optimal *α* is likely approximates 0.5, with *s* potentially around 10^−2^. Further analysis was conducted by sampling 1,000 thalamic data points to compute pair-wise distances for segmentation. Despite using a spatial weight of 0.25 and an SH weight of 0.75, which yielded a DSC close 0.8, the disproportional contribution of spatial and SH-derived distances to the total feature distance led to inconsistency in the clustering algorithm’s performance.

**Figure 2:**
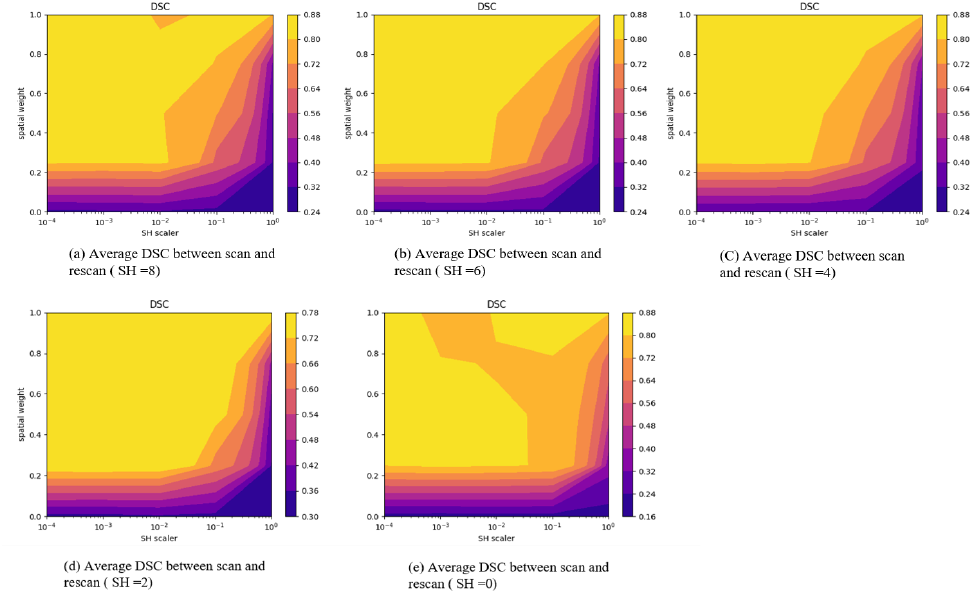
Contour plots illustrating the impact of the spherical harmonic (SH) scaler on the average DSC for 30 scan-rescan subjects in relation to spatial weight, across different SH basis orders. The five plots display a range of SH basis orders: 0, 2, 4, 6, and 8. The X-axis represents the SH coefficient across a spectrum of values: 10^−4^, 10^−3^, 10^−2^, 10^−1^, 10^0^. Y-axis denotes the spatial features weight, with the complementary weight of the SH coefficients calculated as (1 - spatial feature weight). The color gradients indicate the DSC values, with warmer colors denoting higher DSC values.

**Figure 3:**
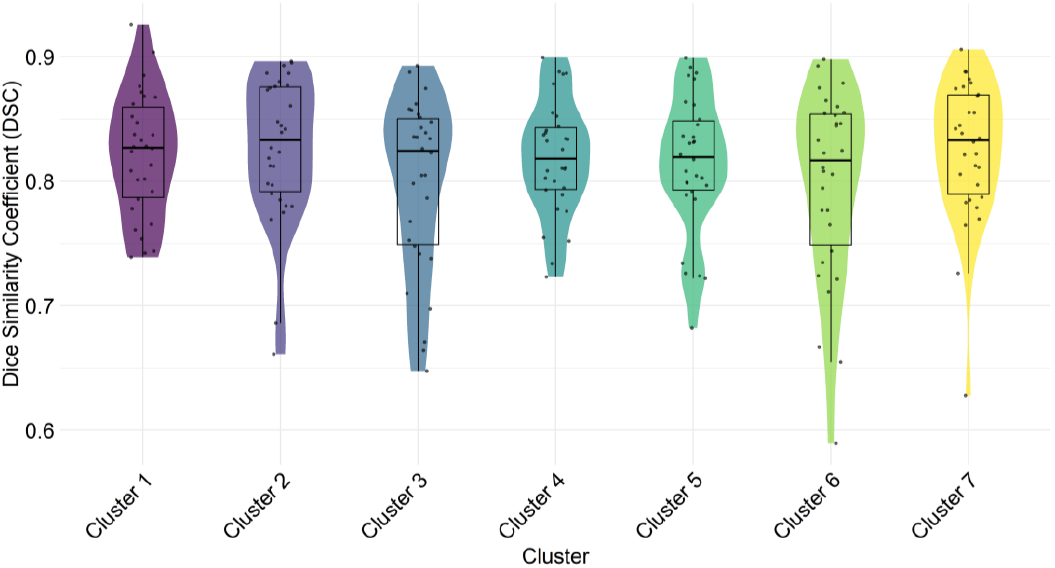
Violin plot of DSC values for 30 test-retest subjects across seven clusters. This violin boxplot visualizes the distribution of Dice Similarity Coefficient (DSC) values across seven clusters for 30 subjects in a test-retest reliability analysis. The X-axis denotes the different clusters (Cluster 1 to Cluster 7), while the Y-axis represents the DSC values. The width of each violin plot indicates the density of DSC values within each cluster, with broader sections representing a higher density of data points. The mean DSC value for each cluster is marked by a black dashed line. Individual data points, plotted as dots within each violin, represent the DSC value for each subject from Dataset A. The DSC values on the Y-axis measure the overlap of specific cluster or nucleus volumes between scan and re-scan sessions, providing an indicator of clustering reliability and consistency.

Notably, equal contribution of spatial and SH-derived distance led to a stable DSC of approximately 0.7. Top 10 DSC values of hyperparameter combinations of BO, with maximum spatial weights of 0.8 (left) and 0.9 (right) provided **in Supplementary Table 1**.

#### 3.1.3. Test-retest reproducibility

The results indicate high within-subject reproducibility, with most DSC values exceeding 0.75. The overall mean DSC was 0.811 (range: 0.798 to 0.826). Clusters with DSCs approaching 0.9 are particularly distinct and well-defined, highlighting the effectiveness of our methodology. Detailed DSC values for each cluster per subject can be found in **Supplementary Table 2**.

### 3.2. Test-retest reproducibility

We assessed the anatomical correspondence between clusters identified by our clustering algorithm and atlas-based thalamic nuclei groups using Dataset A (**Figure 4**). Three clusters showed significant anatomical alignment, Cluster 2 with the Mediodorsal and Central (MDC) group (79.76%), Cluster 6 with the Pulvinar and Geniculate (PuG) group (70.49%), and Cluster 4 with the Ventrolateral and Ventromedial (VLM) group (50.12%). Cluster 3 demonstrated overlaps with multiple nuclei, indicating microstructural similarities, especially with the MDC (27.89%) and Ventral Anterior (VA) (34.23%), and to a lesser extent with VLM (15.75%) and Ventral Posterior (VP) (17.33%). Cluster 5 showed overlaps with VLM (35.37%) and VP (32.86%). Cluster 1’s negligible volume suggests minimal anatomical correspondence. Detailed DSC values for each cluster per subject can be found in **Supplementary Table 3**.

**Figure 4:**
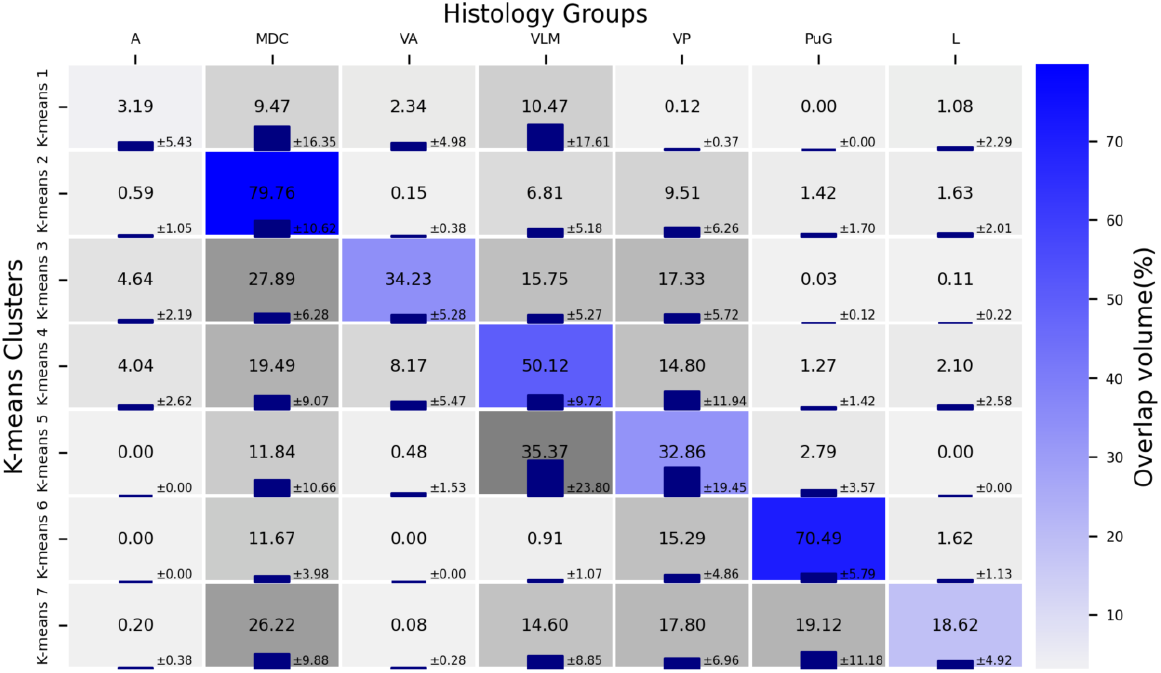
Anatomical correspondence between the FOD based thalamus clusters and atlas-based thalamic nuclei groups. The top X-axis labels seven atlas-based nuclei: the Anterior group (A), Mediodorsal and Central group (MDC), Ventral Anterior group (VA), Ventrolateral and Ventromedial group (VLM), Ventral Posterior group (VP), Pulvinar and Geniculate Nuclei group (PuG), and Lateral groups (L). The Y-axis denotes seven clusters identified by K-means. Each cell displays the mean volume overlap, with the blue error bars representing the standard deviation. The brightness correlates with the mean overlap percentage, reflecting the distribution and variability across the 30 subjects studied (blue code: matrix diagonal, i.e. self-correlation).

### 3.3. Characterizing developmental trajectories of thalamic topography via GMM clustering

Subjects within the dHCP dataset included preterm born infants (n=48) and term born infants (n=68) for analysis.

For the preterm infants in dHCP cohort in Dataset B, significant volumetric increases in all clusters except Cluster 5, with the most substantial growth in Cluster 3 (corresponding to Mediodorsal and Central group (MDC), p < 0.001, pFDR < 0.001, *adjusted* R^2^ = 0.407). Term dHCP infants exhibited notable growth in most clusters, with Cluster 3 (MDC, p < 0.001, pFDR < 0.001, *adjusted* R^2^ = 0.432) and Cluster 5 (Ventral Posterior group (VP), p = 0.001, pFDR = 0.003, *adjusted* R^2^ = 0.219) shown particularly significant increases. The Zurich newborn cohort in Data C presented less pronounced growth, with significant increases only in Clusters 4 (Ventrolateral and Ventromedial group (VLM), p = 0.041, pFDR = 0.076, *adjusted* R^2^ = 0.059) and Cluster 5 show significant increases (Ventral Posterior group (VP), p = 0.001, pFDR = 0.003, *adjusted* R^2^ = -0.042), but only five only out of forty-one subject identified Cluster 5 in this cohort. When analyzing the pooled dataset from all cohorts, significant growth was evident across all clusters, most markedly in Clusters 1 (A), Clusters 2 (MDC), Clusters 3 (VA), Clusters 5 (VP), and Clusters 7 (L).

Our relative volume analysis, which might better reflect topographical changes, did not show statistically significant growth changes in the preterm and term dHCP cohorts in Dataset B. In the Zurich cohort in Dataset C, Cluster 3, corresponding to Ventral Anterior group (VA), suggested a borderline significant increase (p = 0.073). The pooled dataset analysis highlighted significant volume changes in certain clusters, with Cluster 2 showing a notable increase (Mediodorsal and Central group (MDC), p = 0.006, pFDR = 0.021, *adjusted* R^2^ = 0.719), and Cluster 7 (Lateral group (L), p = 0.002, pFDR = 0.012, *adjusted* R^2^ = 0.065).

In the only fetal (post mortem) time-point we tested, the absolute volumes of the clusters were smaller than those of older subjects, such as preterm and term-born infants, consistent with the growth trend of the thalamus. The relative volume proportions showed that the fetal thalamus (at 21 weeks gestational age) exhibited a similar topographical pattern to both preterm and term dHCP infants, with six out of seven clusters having comparable relative volumes. The exception was Cluster 1, which had a volume of zero in the fetal sample, likely due to its early developmental stage with quite small volume (**Figure 8**). Cluster 5 (VP) appeared to increase from fetal to adult age, while the relative proportion of the Cluster 6 (PuG) and Cluster 7 (L) decreased from the mid-gestational fetal time point until newborn age.

Additionally, comparisons with Zurich neonatal and adult data revealed similar relative volumes, suggesting the topographical organization of the thalamus is relatively stable from mid-gestation onwards, however, inter-scanner differences (e.g. between Zurich and dHCP based datasets) appeared to be significant.

#### Statistical analysis of thalamic nuclei volume differences across three cohorts

A more detailed summary of group differences in volume and growth rates see in **Supplementary Table 4**. For the volume difference relative to term-born dHCP infants as the reference group, Cluster 3 in the preterm dHCP cohort showed a significant increase (Ventral Anterior group (VA), standardized β Coef = 1716.11, p = 0.013), with an even more substantial increase observed in the Zurich cohort (standardized β Coef = 3645.36, p < 0.001).

**Table 1.** Absolute volume: robust regression statistics for three datasets, preterm dHCP, term dHCP, and Zurich neonates.

|  |  | Cluster 1 | Cluster 2 | Cluster 3 | Cluster 4 | Cluster 5 | Cluster 6 | Cluster 7 |
| --- | --- | --- | --- | --- | --- | --- | --- | --- |
| **Cohort** |  |  |  |  |  |  |  |  |
| Preterm dHCP | N | 34 | 48 | 48 | 48 | 21 | 41 | 44 |
|  | p | 0.001 | <0.001 | <0.001 | <0.001 | 0.082 | <0.001 | 0.001 |
|  | pFDR | 0.003* | <0.001** | <0.001** | 0.001* | 0.135 | <0.001 | 0.003* |
|  | Ad_R | 0.29 | 0.223 | 0.407 | 0.203 | 0.113 | 0.333 | 0.222 |
| Term dHCP | N | 44 | 68 | 68 | 68 | 32 | 68 | 63 |
|  | p | 0.562 | <0.001 | <0.001 | 0.002 | 0.001 | <0.001 | 0.029 |
|  | pFDR | 0.655 | <0.001** | <0.001** | 0.004* | 0.003* | <0.001** | 0.057 |
|  | Ad_R | -0.02 | 0.147 | 0.432 | 0.120 | 0.219 | 0.281 | 0.069 |
| Zurich Neonates | N | 33 | 27 | 41 | 41 | 5 | 41 | 41 |
|  | p | 0.189 | 0.699 | 0.441 | 0.041 | 0.001 | 0.064 | 0.487 |
|  | pFDR | 0.278 | 0.783 | 0.562 | 0.076 | 0.003* | 0.111 | 0.593 |
|  | Ad_R | 0.029 | -0.060 | -0.024 | 0.059 | -0.042 | 0.049 | 0.002 |
| Pooled dataset | N | 111 | 143 | 157 | 157 | 58 | 157 | 148 |
|  | p | <0.001 | <0.001 | <0.001 | <0.001 | <0.001 | <0.001 | <0.001 |
|  | pFDR | <0.001** | <0.001** | <0.001** | <0.001** | <0.001** | <0.001** | <0.001** |
|  | Ad_R | 0.862 | 0.763 | 0.789 | 0.629 | 0.935 | 0.09 | 0.334 |

**Note:** FDR-corrected significant *p*-values are marked with an asterisk ( $p < 0.05$ ) and a double asterisk ( $p < 0.001$ ); *N*, number of cases; *p*, *p*-value; *pFDR*, FDR corrected *p* value; *Ad\_R*, adjusted *R*-squared value.

**Table 2.** Relative volume robust regression statistics across the preterm dHCP, term dHCP, and Zurich neonatal datasets.

|  |  | Cluster 1 | Cluster 2 | Cluster 3 | Cluster 4 | Cluster 5 | Cluster 6 | Cluster 7 |
| --- | --- | --- | --- | --- | --- | --- | --- | --- |
| **Cohort** |  |  |  |  |  |  |  |  |
| Preterm dHCP | N | 34 | 48 | 48 | 48 | 21 | 41 | 44 |
|  | p | 0.117 | 0.776 | 0.398 | 0.334 | 0.609 | 0.130 | 0.767 |
|  | pFDR | 0.233 | 0.776 | 0.398 | 0.334 | 0.959 | 0.130 | 0.767 |
|  | Ad_R | 0.036 | -0.022 | -0.018 | -0.021 | -0.036 | 0.000 | -0.02 |
| Term dHCP | N | 44 | 68 | 68 | 68 | 32 | 68 | 63 |
|  | p | 0.121 | 0.348 | 0.283 | 0.820 | 0.536 | 0.894 | 0.458 |
|  | pFDR | 0.121 | 0.348 | 0.283 | 0.820 | 0.933 | 0.894 | 0.458 |
|  | Ad_R | 0.032 | -0.006 | 0.003 | -0.014 | -0.003 | -0.015 | -0.011 |
| Zurich Neonates | N | 33 | 27 | 41 | 41 | 5 | 41 | 41 |
|  | p | 0.649 | 0.247 | 0.073 | 0.380 | 0.103 | 0.811 | 0.400 |
|  | pFDR | 0.838 | 0.247 | 0.073 | 0.758 | 0.103 | 0.811 | 0.400 |
|  | Ad_R | -0.024 | -0.016 | 0.060 | 0.003 | 0.263 | -0.024 | -0.015 |
| Pooled dataset | N | 111 | 143 | 157 | 157 | 58 | 157 | 148 |
|  | p | 0.480 | 0.006 | 0.109 | 0.698 | 0.463 | 0.0251 | 0.002 |
|  | pFDR | 0.560 | 0.021* | 0.254 | 0.698 | 0.560 | 0.438 | 0.012* |
|  | Ad_R | 0.854 | 0.719 | 0.854 | 0.557 | 0.892 | 0.260 | 0.065 |

**Note:** FDR-corrected significant *p*-values are marked with an asterisk ( $p < 0.05$ ) and a double asterisk ( $p < 0.001$ ); *N*, Number of cases; *p*, *p*-value; *pFDR*, FDR corrected *p* value; *Ad\_R*, Adjusted *R*-squared value.

**Table 3.** ANOVA test results for absolute volume differences in thalamic nuclei between CHD infants and normal controls.

| Cluster | p-values | pFDR | Effect Size | Power |
| --- | --- | --- | --- | --- |
| Cluster 1 (A) | 0.018 | 0.025* | 0.059 | 0.087 |
| Cluster 2 (MDC) | 0.047 | 0.055 | 0.044 | 0.070 |
| Cluster 3 (VA) | 0.001 | 0.002* | 0.124 | 0.216 |
| Cluster 4 (VLM) | 0.0001 | 0.0004** | 0.150 | 0.292 |
| Cluster 5 (VP) | 0.7182 | 0.718 | 0.002 | 0.050 |
| Cluster 6 (PuG) | 0.004 | 0.007* | 0.086 | 0.128 |
| Cluster 7 (L) | 0.0001 | 0.0004* | 0.154 | 0.307 |

**Note:** FDR-corrected significant p-values are marked with an asterisk ( $p < 0.05$ ) and a double asterisk ( $p < 0.001$ ). N, number of cases; p, p-value; pFDR, FDR corrected p value; Ad\_R, adjusted R-squared value.

**Table 4.**
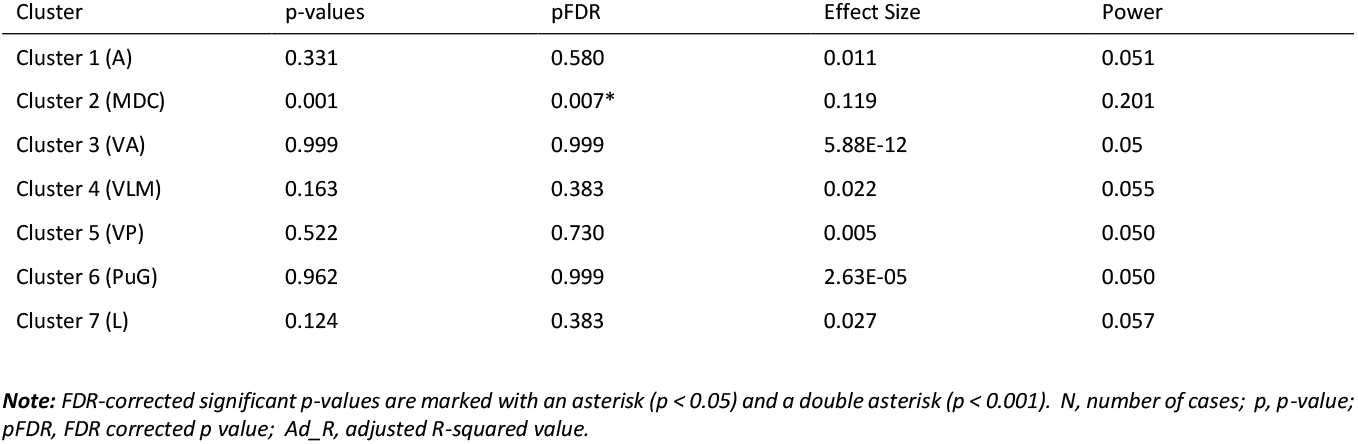
ANOVA test results for relative volume proportion differences in thalamic nuclei between CHD infants and normal controls.

| Cluster | p-values | pFDR | Effect Size | Power |
| --- | --- | --- | --- | --- |
| Cluster 1 (A) | 0.331 | 0.580 | 0.011 | 0.051 |
| Cluster 2 (MDC) | 0.001 | 0.007* | 0.119 | 0.201 |
| Cluster 3 (VA) | 0.999 | 0.999 | 5.88E-12 | 0.05 |
| Cluster 4 (VLM) | 0.163 | 0.383 | 0.022 | 0.055 |
| Cluster 5 (VP) | 0.522 | 0.730 | 0.005 | 0.050 |
| Cluster 6 (PuG) | 0.962 | 0.999 | 2.63E-05 | 0.050 |
| Cluster 7 (L) | 0.124 | 0.383 | 0.027 | 0.057 |

**Note:** FDR-corrected significant p-values are marked with an asterisk ( $p < 0.05$ ) and a double asterisk ( $p < 0.001$ ). N, number of cases; p, p-value; pFDR, FDR corrected p value; Ad\_R, adjusted R-squared value.

### 3.3. Effect of congenital heart disease on thalamus topography

No significant birth age differences were found between the groups. Adjusting for post-menstrual age at MRI, sex, our findings highlighted significantly lower absolute thalamus cluster volumes in six of the seven thalamic nuclei in CHD infants compared to controls. Significantly lower volumes were observed in the Cluster 1 (A), Cluster 3 (VA), the Cluster 4 (VLM), the Cluster 6 (PuG), and the Cluster 7 (L) nuclei groups among CHD infants, with the cluster2 (MDC) group also showing a statistical significant reduction post-correction (pFDR = 0.055). The Cluster 5 (VP) group displayed no significant volume change. Effect sizes were modest for the Cluster 1 (A) and Cluster 2 (MDC) nuclei (0.059 and 0.044, respectively) with significant p-values pre- and post-adjustment. The clusters corresponding to VA, VLM, and PuG nuclei exhibited larger effect sizes (0.124, 0.150, and 0.086, respectively), all maintaining statistical significance after adjustment.

Relative volume analysis showed a significant increase only in the cluster2 (MDC) among CHD infants (p = 0.001, adjusted p = 0.007), indicating an altered topography characterized by this increase. Other nuclei did not exhibit significant relative volume differences between CHD and control groups.

## 4. Discussion

Clustering the thalamus based on MRI features remains an elusive problem that requires optimization and tests of reproducibility and anatomical accuracy. Our study further validated a segmentation framework for clustering thalami nuclei using data from diffusion MRI, leveraging local diffusion properties and spatial features. Our approach is slightly different from existing methods (Mang et al. 2012; Kumar et al. 2015; Battistella et al. 2017), which either utilized lower angular resolution of DWI or applied an arbitrary weighting to the two features. Our approach incorporated white matter FODs from diffusion MRI. Our findings indicate that the consistency of thalamic nuclei segmentation is largely unaffected by whether only the primary eigen-vector or the complete full 45-dimensional set is used, aligning with previous studies that employed only the primary eigenvector from diffusion tensor MRI (Mang et al. 2012; Kumar et al. 2015). Using only the local diffusion characteristics diminished reproducibility in intra-subject thalamic parcellation. This was markedly improved by integrating voxel-wise spatial coordinates, emphasizing the essential role of spatial information in achieving reliable thalamic segmentation. These results highlight the primary eigenvector’s role in reliable thalamic segmentation and suggest that acquiring higher-angular resolution diffusion MRI data may not be necessary for effective dMRI-based thalamic parcellation in such clinically viable scans that we used in our study. We then applied the segmentation method to cluster developing thalami across the perinatal period, scanner, premature born, and in infants with congenital heart defects.

The reproducibility of our thalamic segmentation approach indicated consistent spatial volume overlap in a test-retest analysis across 30 subjects. Applied to multi-centric datasets—including the dHCP, Zurich newborn data, and a human postmortem sample—our method showed comparable volumetric results and proportions of nuclei relative to the total value, which implies a relatively stable thalamus topography.

Our comparisons with the atlas-based mapping showed good spatial alignment for the medial dorsal and central nuclei groups, as well as the pulvinar and geniculate nuclei group. K-means segmented Cluster 4 and Cluster 3 demonstrated overlaps with several atlas-based nuclei. The discrepancies in overlap for some nuclei compared to the transformed Morel atlas might be attributed to two main factors. Firstly, the application of Hausdorff distance in assigning the seven k-means clusters might have led to the overshadowing of smaller nuclei by larger ones, as the latter tend to align more closely with their atlas-defined counterparts. Secondly, methodological differences contribute to these discrepancies; while the Morel atlas delineates nuclei based on cytoarchitectural criteria, our dMRI-based clusters are mostly likely caused by local, dominant fiber structural arrangement. This suggests that, although differences exist, dMRI-based clustering offers a reliable method for identifying thalamic subdivisions, suggesting the technique’s utility in mapping the thalamus’s complex structure.

We evaluated the within-subject reproducibility and anatomical relevance of thalamic segmentation against a seven-cluster grouping of nuclei taken from the Morel atlas (Morel et al. 1997). To identify the optimal number of thalamic clusters, we employed three statistic methods, the elbow method, silhouette score analysis, and gap statistics. These analyses yielded ambiguous results. The elbow method generally indicated 6 or 7 optimal clusters, while both the silhouette score and gap statistics consistently pointed to a binary subdivision as most representative, a finding that differs from Kumar et al. (2015), who consistently identified five clusters across subjects. In our study, although a five-cluster model often corresponded with anatomically meaningful nuclei groups, it was not statistically optimal for thalamic segmentation based on the three statistical methods we used to determine the optimal number of clusters. This multi-method analysis reflects not just the complex structural organization of the thalamus and its developmental trajectory but also underscores the importance of using various methods to fully understand the changes in brain structure during development.

Our study also sheds light on the developmental dynamics of the thalamus, revealing a linear correlation between the absolute volume of thalamic nuclei and scan age (**Figure 5**). This correlation is consistent with biological processes like the maturation of TC connection and the increase in white matter volume (Kostović 2020). Interestingly, relative volumes across the perinatal age, including the premature period, show a stable pattern. This stability aligns with basic research studies suggesting that the topographical organization of the thalamus is established early, following the initial formation of TC circuits and remain stable thereafter (Caviness et al. 1980; Molnár et al. 1998; Kostović and Jovanov-Milošević 2006; Kostovićand Judaš 2010). This concept is encapsulated in the *handshake hypothesis* (Molnar and Blakemore), which proposed that once these early connections are established—around 18th week of gestation—the overall brain structural topology persist through development. Given that our youngest subjects were scanned at 29 gestational weeks, after the TC connections’ initial formation, it follows that the relative volumes of most nuclei we observed remained stable. The only nuclei we found to be changing the topographical organization in the perinatal period in the combined group analysis were Cluster 2 (Mediodorsal and Central group) and Cluster 7 (Lateral groups). The developmental exuberance of neurons in the MD is known and pronounced in the human brain (Abitz et al. 2007), which might be reflected by our finding that the relative volume of this thalamic cluster decreases with gestational age (**Figure 6**).

**Figure 5:**
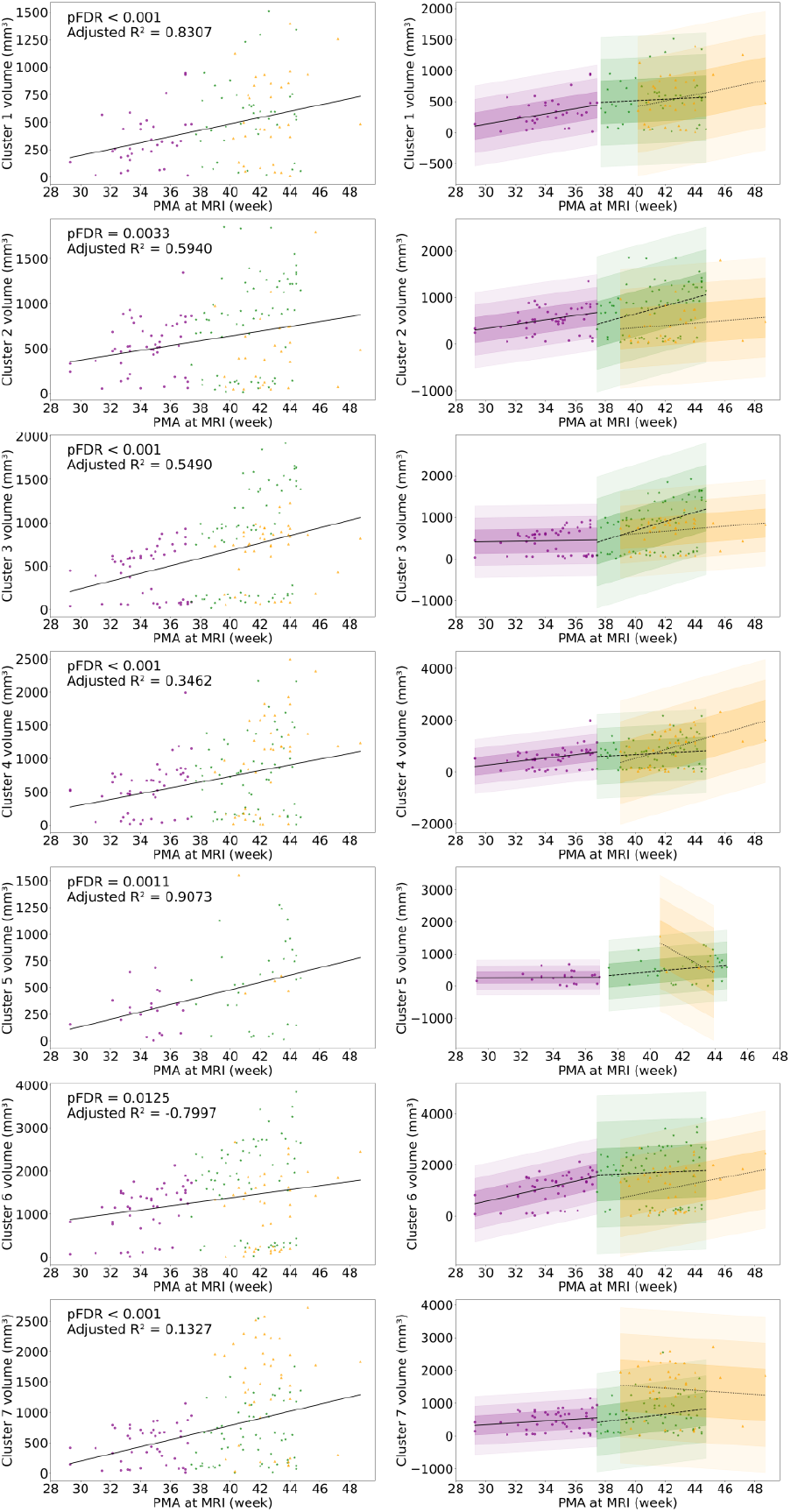
Developmental trajectories of thalamic cluster absolute volumes. The left panel displays the collective regression line generated from combined data across three cohorts, with dots in three distinct colors representing the preterm dHCP, term dHCP, and Zurich neonate cohorts, respectively. The right panel contrasts the regression lines for each cohort separately, with shaded areas representing the variance within ±1, ±2, and ±3 standard deviations, enabling a comparison of their specific growth trajectories.

**Figure 6:**
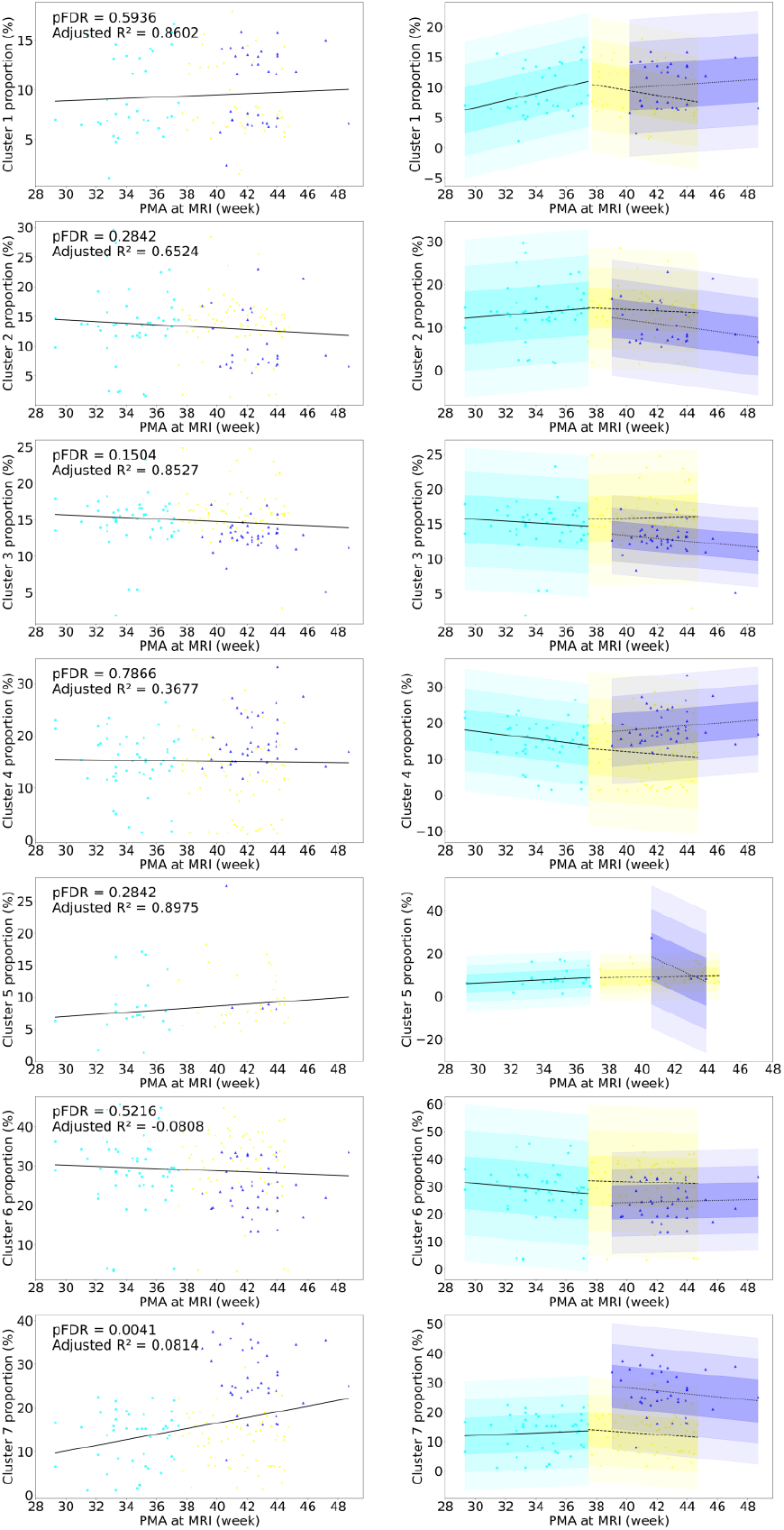
Developmental trajectories of thalamic relative volume. The left panel shows the thalamic volume’s collective regression line based on pooled data from three cohorts, with dots in three different colors representing the preterm dHCP, term dHCP, and Zurich neonate cohorts, respectively. The right panel contrasts these findings with separate cohort-specific regression lines, with shaded areas indicating variance within ±1, ±2, and ±3 standard deviations.

The relative volumes of thalamic clusters indicate that thalamus topography remains stable throughout the perinatal period, as evidenced by similar relative volumes in preterm, term-born infants, Zurich neonates, and adults. Particularly, preterm and term-born infants exhibit nearly identical thalamic nuclei proportions, suggesting that prematurity does not significantly affect thalamic organization. However, significant changes are observed in neonates with congenital heart disease (CHD), notably in Cluster 2 (MDC), which shows an increased relative volume compared to Zurich neonates. Comparing the fetal thalamus to postnatal data reveals early similarities in relative volumes across cohorts. These findings suggest that while the thalamus establishes a stable topographical organization early on, it continues to develop and reorganize, with CHD significantly impacting its structure (**Figure 7)**

**Figure 7.**
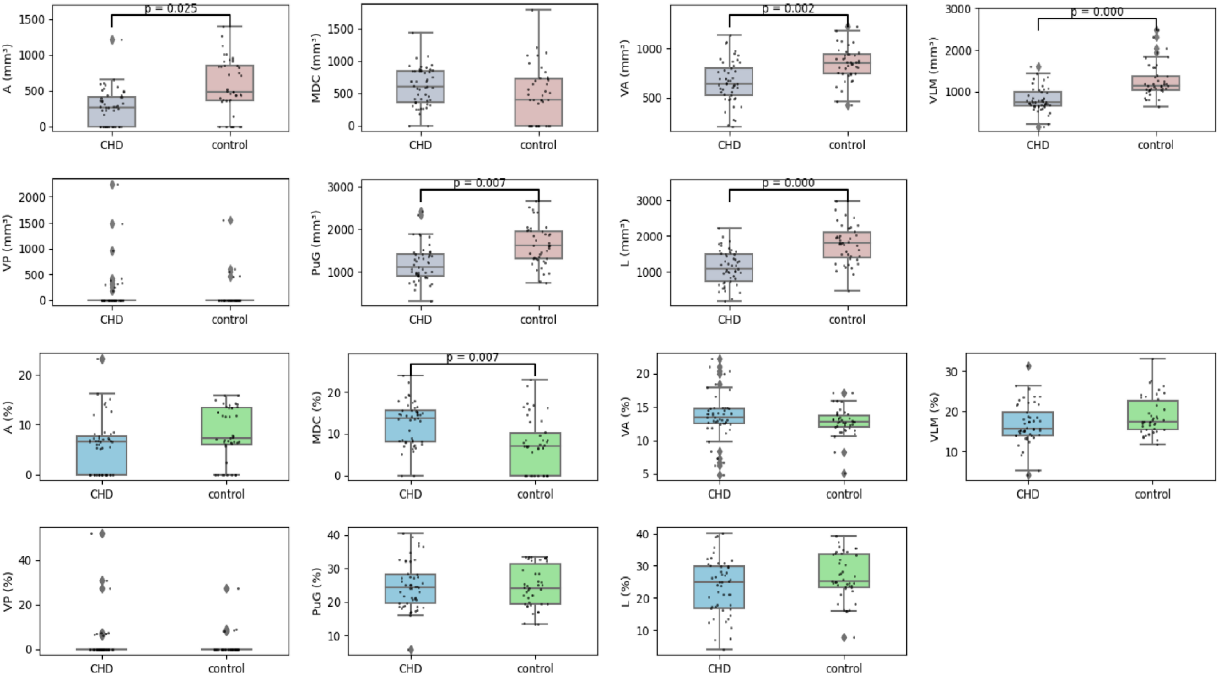
Comparison of thalamic volumes between CHD and control groups. The first and second rows show absolute volume comparisons, while the third and fourth rows show relative volume comparisons, using the seven thalamus clusters.

**Figure 8.**
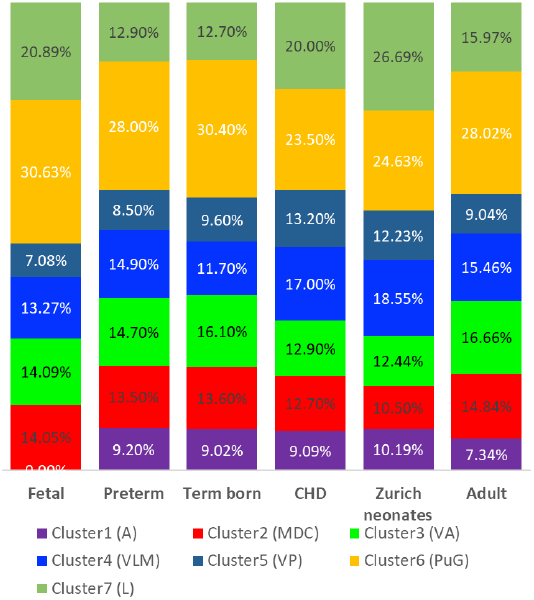
Across-age and across disorder comparison of mean thalamus cluster volumes.

Congenital heart disease (CHD), affecting nearly 1% of live births (Marelli et al. 2016), impairs the delivery of oxygen and nutrients to the brain, contributing to brain injury and increased vulnerability. Our study found a considerable reduction in the volume thalamic nuclei in patients with CHD, consistent with findings of reduced overall brain and sub-cortical volumes, as well as diminished white matter connectivity and delayed cortical development (Von Rhein et al.; Rollins et al.; Limperopoulos et al. 2010; Von Rhein et al. 2015; Rollins et al. 2016). Particularly, the volume of medial dorsal nuclei in CHD infants, relative to the total thalamic volume, points to a targeted impact of CHD on thalamic architecture, potentially affecting whole-brain connectivity and neurodevelopment. Thalamic axons, crucial for the formation neural network and integrity of thalamocortical (TC) projections, play a crucial role in neurodevelopment. Disruptions in TC connectivity, as observed in CHD and preterm births (Gertsvolf et al.; Ball et al. 2012; Ball et al. 2013; Ball et al. 2015; Jaimes et al. 2018), may lead to a reduced thalamic volume and changes in topographical organization, paralleling animal studies that demonstrate significant rearrangements of thalamic axons during development (Molnár et al. 2003). So CHD-related structural and connectivity alterations may affect thalamic organization similarly.

One notable limitation is the reliance on atlas-based transformations. The accuracy of registration potentially impacting the precision of our segmentation and volumetric analysis. Additionally, our use of the Morel atlas derived from adult populations introduces another limitation. The absence of age-specific atlases for infants means that the topological and volumetric nuances associated with different developmental stages may not be fully captured or accurately represented.

In summary, our research further validated a diffusion MRI-based method for subdividing the thalamus, applied to both adult and developmental stages. The stability of thalamic volumes across perinatal ages suggests an early establishment of neural connections, consistent with developmental theories, and a relatively little impact of the on-going developmental processes, such as synaptogenesis, on the thalamic topography. The impact of congenital heart disease on thalamic volume and topography emphasizes the need for further studies to link these characteristics to neurodevelopmental outcomes.

## Acknowledgements

This work is supported by the University Research Priority Program (URPP) “Adaptive Brain Circuits in Development and Learning (AdaBD)” of the University of Zurich. A.J. was supported by the Prof. Max Cloetta Foundation and the EMDO Foundation. H.J. and A.J. were supported by the Vontobel Foundation. Dataset B were provided by the developing Human Connectome Project, KCL-Imperial-Oxford Consortium funded by the European Research Council under the European Union Seventh Framework Programme (FP/2007-2013) / ERC Grant Agreement no. [319456]. We are grateful to the families who generously supported the dHCP trial.

**This manuscript is the author’s original version.**

## Supplementary methods: selecting of the optimal number of clusters

For testing the thalamus topography in the developing cohorts, the number of optimal cluster number is unclear. Since there is no single best approach to determine the optimal number of clusters in a dataset, we used three methods to determine the optimal number of clusters for both the k-means and GMM methods. (1) **Elbow method**: This method plots the unexplained variance against the number of clusters. As the number of cluster increases, the explained variance reduces until a point where the benefit diminishes, forming an ‘elbow’ on the graph. We typically choose the cluster count at this ‘elbow’ as the optimal number. For each subject, the unexplained variance was calculated for cluster numbers ranging from 2 to 15. This provided insights into the proportion of the dataset’s variance that remained unexplained for each clustering solution. A key finding was the identification of a specific cluster number for each subject where the decrease in unexplained variance was no longer substantial, indicating the most suitable number of clusters. (2) **Silhouette Score**: The Silhouette score measures how closely related a thalamus data point is to its cluster compared to other clusters. A higher score suggests well-defined thalamic groupings. For each subject, the optimal number of clusters was determined by identifying the highest average Silhouette score, indicating the clustering configuration where the clusters are most distinct and well-separated.

**(3) Gap statistics:** Gap statistics provide a robust approach to contrast the clustering quality of our observed thalamus data against a baseline of randomly generated reference datasets. This method involves three steps. First, clustering the thalamus data with varying total cluster counts (k = 1, 2, …, K), yielding a set of within-dispersion measures for each k. Second, generating B reference datasets. The B reference datasets are generated by Monte Carlo simulation in accordance to the range of each feature from real data. Thus the reference datasets keep the range of the original features of data but without any cluster structure, and calculating with-dispersion measures *W kb* over a range of cluster counts. Third, calculating the Gap statistic, which quantifies the difference between the log of the observed data’s dispersion and the expected value from the reference dataset. A larger Gap suggests better clustering for that particular k.

Computed the Gap statistic as:

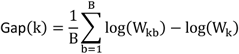

## Supplementary results: optimal number of clusters

We employed three statistical methods to search for the optimal number of clusters in our data: the Elbow Method, Silhouette Score Analysis, and Gap Statistics.

In the test-retest dataset (Data A), the elbow method indicated a preference for 6 or 7 clusters to effectively capture the variance in our data, with 18 subjects favoring 7 clusters and 12 subjects suggesting a better fit with 6. The silhouette score identified 2 clusters as optimal across all 30 subjects. Gap statistics confirmed two clusters as the most common optimal solution. While the majority of subjects (18 out of 30) were optimally clustered into 2 groups, 11 subjects showed a preference for 3 clusters, and one subject uniquely favored 5 clusters.

**Supplementary Figure 1:**
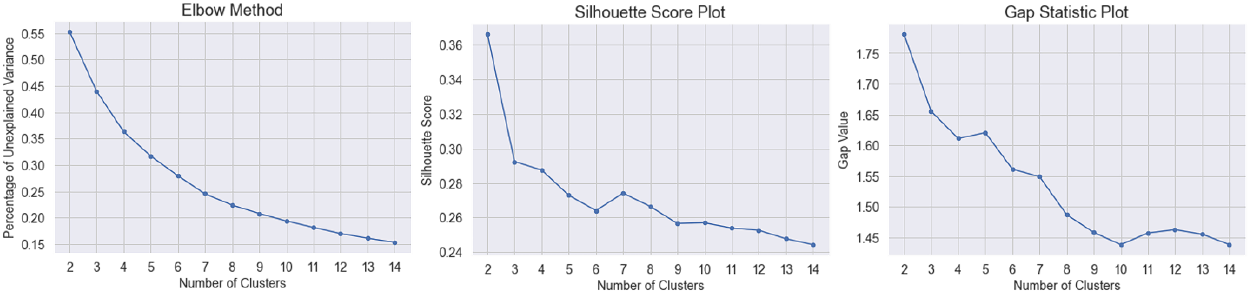
Example of a test-retest subject for comparative analysis of optimal clustering approaches. Left plot shows the Elbow method for a subject from the test-retest data (Data A): the plot displays the mean percentage of unexplained variance against the number of clusters, illustrating a relative clear elbow at 7 clusters. Middle plot shows the silhouette analysis for k-means clustering, the plot shows the average silhouette scores for different numbers of clusters, with the highest score change at 2 clusters. Right side figure shows the gap statistic plot: depicting the gap statistic values across a range of cluster numbers, with the optimal number of clusters indicated by the highest gap value at 2 clusters, suggesting that two clusters provide the most significant statistical differentiation from an expected null reference distribution of the data.

In the dHCP data (Dataset B), the elbow analysis identified six clusters as optimal for explaining data variance in 111 cases, indicating minimal gains with additional clusters. In five instances, seven clusters proved optimal. The silhouette score consistently favored a two-cluster solution as most distinct for our dataset. Gap statistics showed variable optimal cluster numbers. Most subjects were best represented by a two-cluster model, aligning with earlier gestational ages, while a subset, particularly those at later gestational stages (32 weeks and beyond), showed higher optimal cluster numbers. These findings, coupled with analyses of neonatal datasets from Zurich and CHD cases, demonstrate a binary subdivision in earlier developmental stages and increased clustering complexity in later stages.

**Supplementary Table 1:** Bayesian optimization of 100 feature parameters based on average DSC across 30 subjects.

| DSCs | Log SH coefficient | Spatial weight | DSCs | log_SH_coefficient | spatial_weight |
| --- | --- | --- | --- | --- | --- |
| 0.813767733 | -2.933755026 | 0.432840102 | 0.804132132 | -2.808789318 | 0.653510631 |
| 0.813739369 | -2.999547218 | 0.361028328 | 0.804078573 | -1.652061189 | 0.640604768 |
| 0.812198283 | -2.59344781 | 0.221454732 | 0.804034456 | -2.694937791 | 0.168812718 |
| 0.811852681 | -1.868079676 | 0.72162609 | 0.803858106 | -2.703007809 | 0.705479152 |
| 0.811444767 | -4 | 0.553048758 | 0.803747031 | -2.632953376 | 0.517016823 |
| 0.810940296 | -2.755852594 | 0.364549219 | 0.803745141 | -2.460251182 | 0.898054584 |
| 0.810725677 | -2.13162033 | 0.366334117 | 0.803457383 | -1.695036539 | 0.728773066 |
| 0.810724613 | -1.325504883 | 0.9 | 0.803453938 | -2.719425177 | 0.456666089 |
| 0.810490326 | -2.663600215 | 0.303684051 | 0.803349549 | -2.756645682 | 0.072429666 |
| 0.810271065 | -1.779546047 | 0.727087283 | 0.803271955 | -2.596770524 | 0.635011321 |
| 0.809702453 | -2.773317387 | 0.259979974 | 0.802925388 | -2.88865815 | 0.557505752 |
| 0.808953366 | -3.287341985 | 0.0711538 | 0.802825897 | -1.692787509 | 0.439181906 |
| 0.808911237 | -2.824868345 | 0.466120947 | 0.802816477 | -2.479045015 | 0.216547358 |
| 0.808801378 | -2.47739538 | 0.801943376 | 0.802536791 | -2.884420567 | 0.778098794 |
| 0.808733508 | -2.333823595 | 0.611501644 | 0.801846422 | -1.914889976 | 0.648292044 |
| 0.808654306 | -2.663740311 | 0.028294045 | 0.801699318 | -2.376035956 | 0.73125607 |
| 0.80847501 | -2.54395423 | 0.313307911 | 0.801633161 | -1.63467291 | 0.527106605 |
| 0.808019933 | -3.101796234 | 0.369580578 | 0.801454004 | -3.018238555 | 0.557732949 |
| 0.807910795 | -3.171700065 | 0.292969145 | 0.801083557 | -3.331870876 | 0.756855737 |
| 0.807740975 | -2.464155901 | 0.106863173 | 0.8003689 | -2.930060192 | 0.272920096 |
| 0.807671799 | -3.264339578 | 0.089916263 | 0.799997757 | -3.008287472 | 0.898935466 |
| 0.807671796 | -2.603186193 | 0.391997833 | 0.799899434 | -1.780928136 | 0.888397059 |
| 0.80766989 | -2.478759267 | 0.700981874 | 0.799819889 | -2.032013567 | 0.059278978 |
| 0.807586171 | -3.268755323 | 0.085552129 | 0.799617001 | -1.037266736 | 0.782373281 |
| 0.807547218 | -3.018152621 | 0.173842833 | 0.799337745 | -2.283609786 | 0.575355864 |
| 0.807493538 | -2.586649975 | 0.748187853 | 0.799279737 | -1.869075067 | 0.575542548 |
| 0.807457924 | -3.061262182 | 0.272891411 | 0.799142931 | -1.772629912 | 0.479455605 |
| 0.807361315 | -1.830249138 | 0.798493761 | 0.799034317 | -3.175588475 | 0.454974269 |
| 0.80715097 | -2.555835036 | 0.85507682 | 0.799018486 | -2.516395784 | 0.451008525 |
| 0.807146971 | -1.745396389 | 0.804428033 | 0.798803732 | -3.970927982 | 0.9 |
| 0.807026799 | -1.876537335 | 0.886833087 | 0.798571403 | -1.826859255 | 0.650532559 |
| 0.806996509 | -2.87129883 | 0.361710554 | 0.798551902 | -2.607831717 | 0.108289108 |
| 0.806728764 | -1.917219125 | 0.785432944 | 0.798287181 | -2.537135744 | 0.898027483 |
| 0.806480253 | -1.911002711 | 0.64842533 | 0.797556383 | -1.079714737 | 0.778816298 |
| 0.806173634 | -3.266220546 | 0.083104735 | 0.797299147 | -2.902581953 | 0.749543616 |
| 0.806149808 | -2.544693749 | 0.010114717 | 0.797169277 | -2.492877053 | 0.574587092 |
| 0.806048858 | -3.030828327 | 0.450055669 | 0.797150275 | -3.679423062 | 0.702877908 |
| 0.805775372 | -2.365045324 | 0.156992919 | 0.796687121 | -1.278883226 | 0.577669758 |
| 0.805686649 | -1.718362474 | 0.563652533 | 0.795243599 | -1.050605963 | 0.726504873 |
| 0.805623435 | -2.38987602 | 0.038964841 | 0.795126838 | -3.253826836 | 0.233492925 |
| 0.805580572 | -1.727204431 | 0.659391173 | 0.793803143 | -1.03675647 | 0.782873815 |
| 0.805499162 | -2.755737504 | 0.562435735 | 0.746586572 | -0.629683174 | 0.867355964 |
| 0.80544722 | -3.281103621 | 0.100151108 | 0.72646716 | -1.188853455 | 0.295147028 |
| 0.805426441 | -2.929032792 | 0.641444626 | 0.714510045 | -1.666693632 | 0.040724334 |
| 0.805364706 | -2.828185158 | 0.162799967 | 0.655111379 | -0.943168955 | 0.446473765 |
| 0.805327602 | -3.142819853 | 0.18421609 | 0.412450827 | 0.144682915 | 0.886519837 |
| 0.805274074 | -2.661145723 | 0.81628902 | 0.389593069 | -0.164234397 | 0.605842514 |
| 0.805241746 | -1.795280872 | 0.397468123 | 0.30656265 | -0.117394977 | 0.176621886 |
| 0.805203456 | -1.87064267 | 0.46689605 | 0.302968278 | 0.438485155 | 0.402480225 |
| 0.805154152 | -2.792509019 | 0.77128795 | 0.301453952 | -3.799035828 | 0 |
| 0.805110872 | -2.392919646 | 0.517685625 |  |  |  |
| 0.805034995 | -3.999428126 | 0.272099315 |  |  |  |
| 0.80420724 | -3.257775776 | 0.366015314 |  |  |  |

**Supplementary Table 2.** Dice coefficient of scan-rescan data for 30 subjects.

| Subject | Cluster 1 | Cluster 2 | Cluster 3 | Cluster 4 | Cluster 5 | Cluster 6 | Cluster 7 | Average |
| --- | --- | --- | --- | --- | --- | --- | --- | --- |
| 1 | 0.7917 | 0.8868 | 0.6973 | 0.7946 | 0.8214 | 0.7109 | 0.8315 | 0.804074347 |
| 2 | 0.739 | 0.6861 | 0.7379 | 0.7519 | 0.724 | 0.7766 | 0.797 | 0.755486429 |
| 3 | 0.8471 | 0.8185 | 0.8746 | 0.8256 | 0.722 | 0.6667 | 0.7647 | 0.815082699 |
| 5 | 0.8683 | 0.8123 | 0.8571 | 0.886 | 0.8916 | 0.8247 | 0.8789 | 0.802522272 |
| 6 | 0.885 | 0.8951 | 0.8512 | 0.8523 | 0.8313 | 0.8544 | 0.8218 | 0.825935513 |
| 7 | 0.8239 | 0.893 | 0.8389 | 0.7231 | 0.8613 | 0.7649 | 0.8819 | 0.837963492 |
| 8 | 0.8717 | 0.798 | 0.8341 | 0.8337 | 0.8305 | 0.8081 | 0.8878 | 0.856939495 |
| 9 | 0.7654 | 0.6611 | 0.6641 | 0.8782 | 0.8044 | 0.7766 | 0.9058 | 0.827161819 |
| 10 | 0.8763 | 0.8479 | 0.7525 | 0.7759 | 0.7989 | 0.8647 | 0.7848 | 0.796693683 |
| 11 | 0.7854 | 0.7803 | 0.8429 | 0.8548 | 0.8175 | 0.8788 | 0.8057 | 0.733297229 |
| 12 | 0.8372 | 0.876 | 0.8354 | 0.8 | 0.6824 | 0.8328 | 0.8111 | 0.808641315 |
| 13 | 0.8675 | 0.8606 | 0.8537 | 0.8084 | 0.7918 | 0.8549 | 0.8688 | 0.82844606 |
| 14 | 0.7437 | 0.7749 | 0.7481 | 0.7546 | 0.8026 | 0.7241 | 0.7829 | 0.795444578 |
| 15 | 0.9037 | 0.8446 | 0.8876 | 0.8108 | 0.8454 | 0.8444 | 0.7697 | 0.845258832 |
| 16 | 0.8328 | 0.79 | 0.8477 | 0.8389 | 0.726 | 0.8927 | 0.8696 | 0.820751637 |
| 17 | 0.8131 | 0.8963 | 0.7416 | 0.8098 | 0.7857 | 0.811 | 0.8743 | 0.804950535 |
| 18 | 0.8012 | 0.8418 | 0.8262 | 0.8438 | 0.7966 | 0.8056 | 0.8344 | 0.771282047 |
| 19 | 0.926 | 0.8235 | 0.858 | 0.7774 | 0.7889 | 0.8462 | 0.8551 | 0.848283529 |
| 20 | 0.828 | 0.8269 | 0.8228 | 0.8323 | 0.8362 | 0.7944 | 0.8448 | 0.813670695 |
| 21 | 0.8018 | 0.7799 | 0.8049 | 0.8878 | 0.8497 | 0.7437 | 0.7872 | 0.806338996 |
| 22 | 0.8262 | 0.88 | 0.798 | 0.8103 | 0.8081 | 0.8529 | 0.8213 | 0.828734308 |
| 23 | 0.8276 | 0.7851 | 0.6477 | 0.7338 | 0.8356 | 0.5893 | 0.628 | 0.775160253 |
| 24 | 0.852 | 0.877 | 0.8047 | 0.8025 | 0.885 | 0.8599 | 0.8421 | 0.829637825 |
| 25 | 0.8376 | 0.8867 | 0.7864 | 0.8408 | 0.8822 | 0.8226 | 0.8385 | 0.830929816 |
| 26 | 0.8622 | 0.8393 | 0.8928 | 0.8994 | 0.887 | 0.875 | 0.8877 | 0.837834448 |
| 27 | 0.7535 | 0.8733 | 0.8357 | 0.8341 | 0.8318 | 0.8978 | 0.8125 | 0.812636226 |
| 30 | 0.7609 | 0.7969 | 0.7097 | 0.7931 | 0.7985 | 0.7213 | 0.7261 | 0.766376615 |
| 31 | 0.8084 | 0.7692 | 0.671 | 0.789 | 0.734 | 0.6545 | 0.7788 | 0.787196428 |
| 33 | 0.7422 | 0.812 | 0.7676 | 0.8373 | 0.8991 | 0.7348 | 0.8551 | 0.819454879 |
| 34 | 0.7778 | 0.875 | 0.8621 | 0.8866 | 0.8636 | 0.8462 | 0.8759 | 0.84613353 |

**Supplementary Table 3.** Dice coefficient overlap (mean (SD) with the Atlas-based thalamus nuclei groups over 30 subjects.

| K-means/Histology | Histology 1 | Histology 2 | Histology 3 | Histology 4 | Histology 5 | Histology 6 | Histology 7 |
| --- | --- | --- | --- | --- | --- | --- | --- |
| K-means 1 | 3.19 (5.43) | 9.47 (16.35) | 2.34 (4.98) | 10.47 (17.61) | 0.12 (0.37) | 0.00 (0.00) | 1.08 (2.29) |
| K-means 2 | 0.59 (1.05) | 79.76 (10.62) | 0.15 (0.38) | 6.81 (5.18) | 9.51 (6.26) | 1.42 (1.70) | 1.63 (2.01) |
| K-means 3 | 4.64 (2.19) | 27.89 (6.28) | 34.23 (5.28) | 15.75 (5.27) | 17.33 (5.72) | 0.03 (0.12) | 0.11 (0.22) |
| K-means 4 | 4.04 (2.62) | 19.49 (9.07) | 8.17 (5.47) | 50.12 (9.72) | 14.80 (11.94) | 1.27 (1.42) | 2.10 (2.58) |
| K-means 5 | 0.00 (0.00) | 11.84 (10.66) | 0.48 (1.53) | 35.37 (23.80) | 32.86 (19.45) | 2.79 (3.57) | 0.00 (0.00) |
| K-means 6 | 0.00 (0.00) | 11.67 (3.98) | 0.00 (0.00) | 0.91 (1.07) | 15.29 (4.86) | 70.49 (5.79) | 1.62 (1.13) |
| K-means 7 | 0.20 (0.38) | 26.22 (9.88) | 0.08 (0.28) | 14.60 (8.85) | 17.80 (6.96) | 19.12 (11.18) | 18.62 (4.92) |

**Supplementary Table 4.** Comparative analysis of thalamic nuclei volume differences and growth Rates across Preterm dHCP, Term dHCP, and Zurich Neonatal cohorts.

| Cluster | Volume Diff - Preterm<br>vs. Term (Coef, p-value) | Volume Diff - Zurich vs.<br>Term (Coef, p-value) | Growth Rate - Term<br>(Coef, p-value) | Growth Rate - Pre-<br>term (Coef, p-value) | Growth Rate - Zur-<br>ich (Coef, p-value) |
| --- | --- | --- | --- | --- | --- |
| 1 | -1628.88 (p=0.152) | -766.21 (p=0.568) | 22.12 (p=0.249) | 44.15 (p=0.147) | 17.27 (p=0.586) |
| 2 | -42.23 (p=0.964) | 1792.24 (p=0.143) | 57.61 (p=0.000) | 2.25 (p=0.929) | -51.11 (p=0.078) |
| 3 | 1716.11 (p=0.013) | 3645.36 (p=0.000) | 102.66 (p=0.000) | -45.34 (p=0.015) | -96.92 (p=0.000) |
| 4 | 290.82 (p=0.816) | 178.28 (p=0.907) | 74.76 (p=0.000) | -3.93 (p=0.907) | -1.40 (p=0.969) |
| 5 | 1215.20 (p=0.374) | 11040.00 (p=0.003) | 68.60 (p=0.001) | -29.62 (p=0.421) | -261.02 (p=0.003) |
| 6 | 1985.61 (p=0.208) | 3116.15 (p=0.104) | 160.72 (p=0.000) | -55.06 (p=0.192) | -94.41 (p=0.038) |
| 7 | -15.46 (p=0.992) | 1210.58 (p=0.486) | 58.66 (p=0.018) | 1.92 (p=0.961) | -11.25 (p=0.785) |

## References

Abitz M, Nielsen RD, Jones EG, Laursen H, Graem N, Pakkenberg B. 2007. Excess of neurons in the human newborn mediodorsal thalamus compared with that of the adult. Cereb Cortex.s 17(11):2573–2578. doi:10.1093/CER-COR/BHL163.

Alcauter S, Lin W, Smith XJK, Short SJ, Goldman BD, Reznick JS, Gilmore JH, Gao W. 2014. Development/Plasticity/Repair Development of Thalamocortical Connectivity during Infancy and Its Cognitive Correlations. doi:10.1523/JNEUROSCI.0796-14.2014.

Alexander B, Kelly CE, Adamson C, Beare R, Zannino D, Chen J, Murray AL, Loh WY, Matthews LG, Warfield SK, et al. 2018. Changes in neonatal regional brain volume associated with preterm birth and perinatal factors. doi:10.1016/j.neuroimage.2018.07.021.

Andersson JLR, Skare S, Ashburner J. 2003. How to correct susceptibility distortions in spin-echo echo-planar images: application to diffusion tensor imaging. Neuroimage. 20(2):870–888. doi:10.1016/S1053-8119(03)00336-7.

Andersson JLR, Sotiropoulos SN. 2016. An integrated approach to correction for off-resonance effects and subject movement in diffusion MR imaging. Neuroimage. 125:1063–1078. doi:10.1016/j.neuroimage.2015.10.019.

Anticevic A, Cole MW, Repovs G, Murray JD, Brumbaugh MS, Winkler AM, Savic A, Krystal JH, Pearlson GD, Glahn DC. 2014. Characterizing Thalamo-Cortical Disturbances in Schizophrenia and Bipolar Illness. Cereb Cortex (New York, NY). 24(12):3116. doi:10.1093/CERCOR/BHT165.

Auladell C, Pérez-Sust P, Supèr H, Soriano E. 2000. The early development of thalamocortical and corticothalamic projections in the mouse. Anat Embryol (Berl). 201(3):169–179. doi:10.1007/PL00008238/METRICS.

Avants BB, Epstein CL, Grossman M, Gee JC. 2008. Symmetric diffeomorphic image registration with cross-correlation: Evaluating automated labeling of elderly and neurodegenerative brain. Med Image Anal. 12(1):26–41. doi:10.1016/J.MEDIA.2007.06.004.

Avants BB, Tustison NJ, Song G, Cook PA, Klein A, Gee JC. 2011. A reproducible evaluation of ANTs similarity metric performance in brain image registration. Our Symmetric Norm NeuroImage. 54:2033–2044. doi:10.1016/j.neuroimage.2010.09.025.

Ball G, Boardman JP, Aljabar P, Pandit A, Arichi T, Merchant N, Rueckert D, Edwards AD, Counsell SJ. 2013. The influence of preterm birth on the developing thalamocortical connectome. Cortex. 49(6):1711–1721. doi:10.1016/J.CORTEX.2012.07.006.

Ball G, Boardman JP, Rueckert D, Aljabar P, Arichi T, Merchant N, Gousias IS, David Edwards A, Counsell SJ, Steiner R. 2012. The Effect of Preterm Birth on Thalamic and Cortical Development. Cereb Cortex. 22:1016–1024. doi:10.1093/cercor/bhr176.

Ball G, Pazderova L, Chew A, Tusor N, Merchant N, Arichi T, Allsop JM, Cowan FM, Edwards AD, Counsell SJ. 2015. Thalamocortical Connectivity Predicts Cognition in Children Born Preterm. Cereb Cortex (New York, NY). 25(11):4310. doi:10.1093/CERCOR/BHU331.

Bastiani M, Andersson JLR, Cordero-Grande L, Murgasova M, Hutter J, Price AN, Makropoulos A, Fitzgibbon SP, Hughes E, Rueckert D, et al. 2018. Automated processing pipeline for neonatal diffusion MRI in the developing Human Connectome Project. doi:10.1016/j.neuroimage.2018.05.064.

Battistella Giovanni, Najdenovska Elena, Maeder P, Naghmeh Ghazaleh •, Daducci A, Thiran J-P, Jacquemont S, Constantin Tuleasca •, Levivier M, Meritxell •, et al. 2017. Robust thalamic nuclei segmentation method based on local diffusion magnetic resonance properties. Brain Struct Funct. 222:2203–2216. doi:10.1007/s00429-016-1336-4.

Behrens TEJ, Johansen-Berg H, Woolrich MW, Smith SM, Wheeler-Kingshott CAM, Boulby PA, Barker GJ, Sillery EL, Sheehan K, Ciccarelli O, et al. 2003. Non-invasive mapping of connections between human thalamus and cortex using diffusion imaging. Nat Neurosci 2003 67. 6(7):750–757. doi:10.1038/nn1075.

Bertholdt S, Latal B, Liamlahi R, Prêtre R, Scheer I, Goetti R, Dave H, Bernet V, Schmitz A, Von Rhein M, et al. 2013. Cerebral lesions on magnetic resonance imaging correlate with preoperative neurological status in neonates undergoing cardiopulmonary bypass surgery. doi:10.1093/ejcts/ezt422.

Boekel W, Forstmann BU, Keuken MC. 2016. A test-retest reliability analysis of diffusion measures of white matter tracts relevant for cognitive control. doi:10.1111/psyp.12769.

Brossard-Racine Gilbert M, Easson K, Majnemer A, Marelli A, Fontes MK, Courtin F, Rohlicek C, Saint-Martin C, Fontes K, Gilbert G, et al. 2020. Characterizing the Subcortical Structures in Characterizing the Subcortical Structures in Youth with Congenital Heart Disease. AJNR Am J Neuroradiol. 41(8):1503–1508. doi:10.3174/ajnr.A6667.

Carrera E, Bogousslavsky J. 2006. The thalamus and behavior Effects of anatomically distinct strokes. Neurology. 66:1817–1823.

Caviness VS, and JR, Frost DO, Caviness S, Ken E. 1980. Organization of Thalamic Projections to the in the Mouse Send reprint requests to: Verne. J Comp Neurol.:194335–367.

Cordero-Grande L, Hughes EJ, Hutter J, Price AN, Hajnal J V. Three-Dimensional Motion Corrected Sensitivity Encoding Reconstruction for Multi-Shot Multi-Slice MRI: Application to Neonatal Brain Imaging. doi:10.1002/mrm.26796.

Dhollander T, Connelly A. 2016. A novel iterative approach to reap the benefits of multi-tissue CSD from just single-shell (+b=0) diffusion MRI data. Proc Intl Soc Mag Reson Med 24.(May).

Dhollander T, Mito R, Raffelt D, Connelly A. 2019. Improved white matter response function estimation for 3-tissue constrained spherical deconvolution. Proc Intl Soc Mag Reson Med.(May 11-16):555.

Dubois J, Dehaene-Lambertz G, Kulikova S, Poupon C, Hüppi PS, Hertz-Pannier L. 2014. The early development of brain white matter: A review of imaging studies in fetuses, newborns and infants. Neuroscience. 276:48–71. doi:10.1016/J.NEUROSCIENCE.2013.12.044.

Eugenio Iglesias J, Insausti R, Lerma-Usabiaga G, Bocchetta M, Van Leemput K, Greve DN, van der Kouwe A, Alzheimer the, Neuroimaging Initiative D, Fischl B, et al. 2018. A probabilistic atlas of the human thalamic nuclei combining ex vivo MRI and histology. doi:10.1016/j.neuroimage.2018.08.012.

Feldmann M, Guo T, Miller SP, Knirsch W, Kottke R, Hagmann C, Latal B, Jakab A. Delayed maturation of the structural brain connectome in neonates with congenital heart disease. doi:10.1093/braincomms/fcaa209.

Gertsvolf N, Votava-Smith JK, Ceschin R, Del Castillo S, Lee V, Lai HA, Bluml S, Paquette L, Panigrahy A. Association between Subcortical Morphology and Cerebral White Matter Energy Metabolism in Neonates with Congenital Heart Disease OPEN. doi:10.1038/s41598-018-32288-3.

Gousias IS, Edwards AD, Rutherford MA, Counsell SJ, Hajnal J V., Rueckert D, Hammers A. 2012. Magnetic resonance imaging of the newborn brain: Manual segmentation of labelled atlases in term-born and preterm infants. Neuroimage. 62(3):1499–1509. doi:10.1016/J.NEUROIMAGE.2012.05.083.

Iglehart C, Monti M, Cain J, Tourdias T, Saranathan M. 2020. A systematic comparison of structural-, structural connectivity-, and functional connectivity-based thalamus parcellation techniques. Brain Struct Funct. 225(5):1631– 1642. doi:10.1007/S00429-020-02085-8/FIGURES/6.

Jaimes C, Cheng HH, Soul J, Ferradal S, Rathi Y, Gagoski B, Newburger JW, Grant PE, Zöllei L. 2018. Probabilistic tractography-based thalamic parcellation in healthy newborns and newborns with congenital heart disease. J Magn Reson Imaging. 47(6):1626–1637. doi:10.1002/JMRI.25875.

Jakab A, Blanc R, Berényi EL, Székely G. 2012. Generation of Individualized Thalamus Target Maps by Using Statistical Shape Models and Thalamocortical Tractography. Am J Neuroradiol. 33(11):2110–2116. doi:10.3174/AJNR.A3140.

Jbabdi S, Woolrich MW, Behrens TEJ. 2009. Multiple-subjects connectivity-based parcellation using hierarchical Dirichlet process mixture models. Neuroimage. 44(2):373–384. doi:10.1016/J.NEUROIMAGE.2008.08.044.

Jenkinson M, Smith S. 2001. A global optimisation method for robust affine registration of brain images. Med Image Anal. 5(2):143–156. doi:10.1016/S1361-8415(01)00036-6.

Jonasson L, Hagmann P, Pollo C, Bresson X, Wilson CR, Meuli R, Thiran J-P. 2007. A level set method for segmentation of the thalamus and its nuclei in DTMRI. Signal Processing. 87:309–321. doi:10.1016/j.sigpro.2005.12.017.

Jones EG. 2007. The Thalamus, Volume 1. Cambridge University Press.

Jorge Cardoso M, Li W, Brown R, Ma N, Kerfoot E, Wang Y, Murrey B, Myronenko A, Zhao C, Yang D, et al. 2022. MONAI: An open-source framework for deep learning in healthcare.

Kikinis R, Pieper SD, Vosburgh KG. 2014. 3D Slicer: A Platform for Subject-Specific Image Analysis, Visualization, and Clinical Support. Intraoperative Imaging Image-Guided Ther.:277–289. doi:10.1007/978-1-4614-7657-3_19.

Kim J, Choi S. 2023. BayesO: A Bayesian optimization framework in Python. J Open Source Softw. 8(90):5320. doi:10.21105/JOSS.05320.

Kostović I. 2020. The enigmatic fetal subplate compartment forms an early tangential cortical nexus and provides the framework for construction of cortical connectivity. doi:10.1016/j.pneurobio.2020.101883.

Kostović I, Jovanov-Milošević N. 2006. The development of cerebral connections during the first 20–45 weeks’ gestation. Semin Fetal Neonatal Med. 11(6):415–422. doi:10.1016/J.SINY.2006.07.001.

Kostović I, Judaš M. 2010. The development of the subplate and thalamocortical connections in the human foetal brain. Acta Paediatr Int J Paediatr. 99(8):1119–1127. doi:10.1111/J.1651-2227.2010.01811.X.

Krauth A, Blanc R, Poveda A, Jeanmonod D, Morel A, Székely G. 2009. A mean three-dimensional atlas of the human thalamus: Generation from multiple histological data. Neuroimage. 49:2053–2062. doi:10.1016/j.neuroimage.2009.10.042.

Kubota M, Miyata J, Sasamoto A, Sugihara G, Yoshida H, Kawada R, Fujimoto S, Tanaka Y, Sawamoto N, Fukuyama H, et al. 2013. Thalamocortical Disconnection in the Orbitofrontal Region Associated With Cortical Thinning in Schizophrenia. JAMA Psychiatry. 70(1):12–21. doi:10.1001/ARCHGENPSYCHIATRY.2012.1023.

Kuklisova-Murgasova M, Quaghebeur G, Rutherford MA, Hajnal J V., Schnabel JA. 2012. Reconstruction of fetal brain MRI with intensity matching and complete outlier removal. Med Image Anal. 16(8):1550. doi:10.1016/J.MEDIA.2012.07.004.

Kumar V, Mang S, Grodd W. 2015. Direct diffusion-based parcellation of the human thalamus. Brain Struct Funct. 220(3):1619–1635. doi:10.1007/S00429-014-0748-2/FIGURES/7.

Limperopoulos C, Tworetzky W, McElhinney DB, Newburger JW, Brown DW, Robertson RL, Guizard N, McGrath E, Geva J, Annese D, et al. 2010. Brain Volume and Metabolism in Fetuses With Congenital Heart Disease. Circulation. 121(1):26–33. doi:10.1161/CIRCULATIONAHA.109.865568.

Makropoulos A, Robinson EC, Schuh A, Wright R, Fitzgibbon S, Bozek J, Counsell SJ, Steinweg J, Vecchiato K, Passerat-Palmbach J, et al. 2018. The Developing Human Connectome Project: a Minimal Processing Pipeline for Neonatal Cortical Surface Reconstruction Europe PMC Funders Group. Neuroimage. 173:88–112. doi:10.1101/125526.

Mang SC, Busza A, Reiterer S, Grodd W, Klose U. 2012. Thalamus Segmentation Based on the Local Diffusion Direction: A Group Study. Magn Reson Med. 67:118–126. doi:10.1002/mrm.22996.

Marelli A, Miller SP, Marino BS, Jefferson AL, Newburger JW. 2016. Brain in congenital heart disease across the lifespan: The cumulative burden of injury. Circulation. 133(20):1951–1962. doi:10.1161/CIRCULATIONAHA.115.019881/FORMAT/EPUB.

Menegaux A, Meng C, b JG, Berndt MT, Hedderich DM, Schmitz-Koep B, Schneider S, Nuttall R, Zimmermann J, Daamen M, et al. 2021. Aberrant corticothalamic structural connectivity in premature-born adults. doi:10.1016/j.cortex.2021.04.009.

Miller B, Chou L, Finlay BL. 1993. The early development of thalamocortical and corticothalarnic projections. J Comp Neurol. 335(1):16–41. doi:10.1002/CNE.903350103.

Molnár Z, Adams R, Blakemore C. 1998. Mechanisms Underlying the Early Establishment of Thalamocortical Connections in the Rat. J Neurosci. 18(15):5723. doi:10.1523/JNEUROSCI.18-15-05723.1998.

Molnár Z, Higashi S, López-Bendito G. 2003. Choreography of Early Thalamocortical Development. Cereb Cortex. 13(6):661–669. doi:10.1093/CERCOR/13.6.661.

Molnhr Z, Blakemore C. How do thalamic axons find their way to the cortex?

Morel A, Magnin M, Jeanmonod D. 1997. Multiarchitectonic and Stereotactic Atlas of the Human Thalamus. J Comp Neurol. 387:588–630. doi:10.1002/(SICI)1096-9861(19971103)387:4<588::AID-CNE8>3.0.CO;2-Z.

Nair A, Treiber JM, Shukla DK, Shih P, Mü R-A. Impaired thalamocortical connectivity in autism spectrum disorder: a study of functional and anatomical connectivity. A J Neurol. doi:10.1093/brain/awt079.

Nosarti C, Woo Nam K, Walshe M, Murray RM, Cuddy M, Rifkin L, Allin MP. 2014. Preterm birth and structural brain alterations in early adulthood. doi:10.1016/j.nicl.2014.08.005.

Pedregosa FABIANPEDREGOSA F, Michel V, Grisel OLIVIERGRISEL O, Blondel M, Prettenhofer P, Weiss R, Vanderplas J, Cournapeau D, Pedregosa F, Varoquaux G, et al. 2011. Scikit-learn: Machine Learning in Python Gaël Varoquaux Bertrand Thirion Vincent Dubourg Alexandre Passos PEDREGOSA, VAROQUAUX, GRAMFORT ET AL. Matthieu Perrot. J Mach Learn Res. 12:2825–2830.

Von Rhein M, Buchmann A, Hagmann C, Dave H, Bernet V, Scheer I, Knirsch W, Latal B. 2015. Severe Congenital Heart Defects Are Associated with Global Reduction of Neonatal Brain Volumes. J Pediatr. 167:1259–63. doi:10.1016/j.jpeds.2015.07.006.

Von Rhein M, Buchmann A, Hagmann C, Huber R, Klaver P, Knirsch W, Latal B. Brain volumes predict neurodevelopment in adolescents after surgery for congenital heart disease. A J Neurol. doi:10.1093/brain/awt322.

Rollins CK, Asaro LA, Akhondi-Asl A, Kussman BD, Rivkin MJ, Bellinger DC, Warfield SK, Wypij D, Newburger JW, Soul JS. 2016. White Matter Volume Predicts Language Development in Congenital Heart Disease. J Pediatr. 181:42-48.e2. doi:10.1016/j.jpeds.2016.09.070.

Rollins CK, Watson CG, Asaro LA, Wypij D, Vajapeyam S, Bellinger DC, Demaso DR, Robertson RL, Newburger JW, Rivkin MJ. White Matter Microstructure and Cognition in Adolescents with Congenital Heart Disease. doi:10.1016/j.jpeds.2014.07.028.

Seabold S, Perktold J. 2010. Statsmodels: Econometric and Statistical Modeling with Python. PROC 9th PYTHON Sci CONF.

Su JH, Thomas FT, Kasoff WS, Tourdias T, Choi EY, Rutt BK, Saranathan M. 2019. Thalamus Optimized Multi Atlas Segmentation (THOMAS): fast, fully automated segmentation of thalamic nuclei from structural MRI. doi:10.1016/j.neuroimage.2019.03.021.

Sur M, Rubenstein JLR. 2005. Patterning and plasticity of the cerebral cortex. Science. 310(5749):805–10. doi:10.1126/science.1112070.

Takahata A, Nakagawa M, Mineura K, Nishimura T, Yamada K, Akazawa K, Yuen S, Goto M, Matsushima S. 2010. MR Imaging of Ventral Thalamic Nuclei. AJNR Am J Neuroradiol. 31(4):732–735. doi:10.3174/ajnr.A1870.

Thompson DK, Matthews LG, Alexander B, Lee KJ, Kelly CE, Adamson CL, Hunt RW, Cheong JLY, Spencer-Smith M, Neil JJ, et al. Tracking regional brain growth up to age 13 in children born term and very preterm. doi:10.1038/s41467-020-14334-9.

Tournier J-D, Calamante F, Gadian DG, Connelly A. 2004. Direct estimation of the fiber orientation density function from diffusion-weighted MRI data using spherical deconvolution. doi:10.1016/j.neuroimage.2004.07.037.

Tournier J-D, Smith R, Raffelt D, Tabbara R, Dhollander T, Pietsch M, Christiaens D, Jeurissen B, Yeh C-H, Connelly A. 2019. MRtrix3: A fast, flexible and open software framework for medical image processing and visualisation. doi:10.1016/j.neuroimage.2019.116137.

Tustison NJ, Avants BB, Cook PA, Zheng Y, Egan A, Yushkevich PA, Gee JC. 2010. N4ITK: Improved N3 bias correction. IEEE Trans Med Imaging. 29(6):1310– 1320. doi:10.1109/TMI.2010.2046908.

Unrath A, Klose U, Grodd W, Ludolph AC, Kassubek J. 2008. Directional colour encoding of the human thalamus by diffusion tensor imaging. doi:10.1016/j.neulet.2008.02.013.

Veraart J, Fieremans E, Novikov DS. 2016. Diffusion MRI noise mapping using random matrix theory. Magn Reson Med. 76(5):1582–1593. doi:10.1002/MRM.26059.

Wassermann D, Descoteaux M, Deriche R. 2008. Diffusion Maps Clustering for Magnetic Resonance Q-Ball Imaging Segmentation. Int J Biomed Imaging. doi:10.1155/2008/526906.

Wiegell MR, Tuch DS, Larsson HBW, Wedeen VJ. 2003. Automatic segmentation of thalamic nuclei from diffusion tensor magnetic resonance imaging. doi:10.1016/S1053-8119(03)00044-2.

Ye C, Bogovic JA, Ying SH, Prince JL. 2013. Parcellation of the Thalamus Using Diffusion Tensor Images and a Multi-object Geometric Deformable Model. Proc SPIE. 8669:866909. doi:10.1117/12.2006119.

Ziyan U, Tuch D, Westin CF. 2006. Segmentation of thalamic nuclei from DTI using spectral clustering. Med Image Comput Comput Assist Interv. 9(Pt 2):807–814. doi:10.1007/11866763_99.

Zou KH, Warfield SK, Bharatha A, Tempany CMC, Kaus MR, Haker SJ, Wells Iii WM, Jolesz FA, Kikinis R. 2004. Statistical Validation of Image Segmentation Quality Based on a Spatial Overlap Index 1 Scientific Reports. Radiol Alliance Heal Serv Res Acad Radiol. 11:178–189. doi:10.1016/S1076-6332(03)00671-8.

